# Region-specific CREB function regulates distinct forms of regret associated with resilience versus susceptibility to chronic stress

**DOI:** 10.1101/2022.01.17.476637

**Authors:** Romain Durand-de Cuttoli, Freddyson J. Martínez-Rivera, Long Li, Angélica Minier-Toribio, Flurin Cathomas, Leanne M. Holt, Farzana Yasmin, Salma O. Elhassa, Jasmine F. Shaikh, Sanjana Ahmed, Scott J. Russo, Eric J. Nestler, Brian M. Sweis

## Abstract

Regret describes recognizing that alternative actions could have led to better outcomes. This can transform into behavioral consequences, altering subsequent valuations, but remains unclear if regret derives from a generalized computation for mistake appraisal or instead is made up of dissociable action-specific processes. Using a novel neuroeconomic decision-making paradigm, we found mice were differentially sensitive to fundamentally distinct types of missed opportunities following exposure to chronic social defeat stress or manipulations of CREB, a key transcription factor implicated in chronic stress action. Bias to make compensatory decisions after rejecting high-value offers (regret type I) was unique to stress-susceptible mice. Bias following the converse operation, accepting low-value offers (regret type II), was enhanced in stress-resilient and absent in stress-susceptible mice. CREB function in either the medial prefrontal cortex or nucleus accumbens was required to suppress regret type I but differentially affected regret type II. We provide insight into how adaptive versus maladaptive stress-response traits may be related to fundamentally distinct forms of counterfactual thinking and could steer psychotherapy for mood disorders such as depression toward unveiling circuit-specific computations through a careful description of decision narrative.

---

Mistakes are an essential component of reinforcement learning (*1*). However, acknowledging the error of one’s own agency separates the more complex experience of regret from the mere disappointment of rewards that do not meet expectations (*2*). Counterfactual thinking, or imagining unselected actions, lies at the core of how regret is processed (*3, 4*). Appreciating that an alternative action could have led to a better outcome serves as the computational basis of regret that can alter mood and may detract from one’s emotional well-being, potentially promoting psychiatric syndromes such as depression (*2*). Consideration of regret can influence subsequent valuations, often in a compensatory manner and even sometimes at the expense of making optimal decisions (*5*). This has been postulated in the psychology and economics literature nonetheless to serve to increase the salience of realized losses, which may be useful for mitigating cognitive dissonance and avoiding future regret (*1, 2, 6, 7*).

Stress-related disorders such as depression are debilitating illnesses in which individuals struggle with severe emotional dysregulation (*8*). It is widely accepted that regret contributes to disease burden and may be linked to symptoms including emotional reactivity and negative rumination (*8-10*). On the other hand, a decreased sensitivity to regret may also appear in depression as individuals often suffer from generalized numbness related to anhedonia, a key symptom of this disorder (*8, 9*). Thus, no aspect of regret appears pathognomonic for depression, with differences seen across subtypes of this heterogeneous syndrome. Furthermore, little is known about the neurobiology of what might make this process maladaptive and what aspects of regret, if any, carry utility that is worth preserving in order to restore healthy emotional processing and adaptive coping. Therefore, a more thorough understanding of the computational underpinnings of regret is needed.

Animal models used for the study of depression and other stress-related disorders have made significant contributions to the field. These include identifying key molecular mediators, such as CREB, which regulates the transcription of stress-sensitive genes that control responses to rewarding and stressful stimuli in a brain-region-specific manner (*11-13*). For instance, CREB activation in the nucleus accumbens increases the display of depressive-like traits in rodent models, whereas CREB activation in the medial prefrontal cortex promotes resilience to stress (*13-15*). Analysis of human postmortem brain tissue supports a similar region-specific role in depression (*13*). While elucidating functions of CREB in mediating differential stress responses has begun to make headway, the role of CREB in decision making, let alone the processing of the negative consequences of one’s choices, is far less understood. Furthermore, animal models have been limited in their ability to capture the complexity of affective processes in decision making observed in human patients with stress-related disorders. To shed light on this translational gap, we combined the well-established chronic social defeat stress model for depression in mice with molecular manipulations and novel approaches in neuroeconomics that have been validated for use in both rodents and humans and only recently demonstrated the behavioral and neurophysiological correlates of regret in rodents (*5, 7, 11, 14, 16*).

How can one ascertain if rodents are capable of experiencing regret? How can such a phenomenon be measured non-verbally? In 2014, Steiner and Redish developed a novel decision-making task, termed “Restaurant Row” for use in rats (*5*). Animals foraged for food rewards while on a limited time budget making accept or reject decisions for offers of varying costs in the form of delays signaled by the pitch of a tone. Choices were supported by behavioral and neurophysiological evidence of “deliberation” during tone presentation suggesting animals were comparing competing alternatives before making informed decisions (*17*). Steiner and Redish found that in the specific situation in which rats inappropriately rejected a short delay offer only to encounter a long delay offer on the next trial did animals physically look back at the previous reward site concurrent with “replay” events in frontal and striatal ensembles representing the previous decision-point location leading to the forgone reward. These events served as correlates of regret-related processes and biased animals to be more likely to overcompensate and accept long delay offers on the second trial compared to sequences in which animals did not make economic violations.

Building off of this work, in 2018 Sweis et al. translated this task for use in mice and examined the sequelae of the converse economic violation: inappropriately accepting long delay offers (*7*). The task design was modified such that mice had an opportunity to correct these putative mistakes within the same trial. In this modified version of Restaurant Row, decisions were physically separated into two distinct stages: tones presented in an offer zone that indicated the delay of the current trial but did not begin to countdown until mice explicitly entered a separate wait zone during which mice could quit at any point. This allowed for a more thorough interrogation of how decisions to enter the wait zone for long delay offers might be erroneous. The authors found that the majority of decisions to enter long delay offers occurred on trials in which mice made ballistic journeys through the offer zone. These events signaled a failure to “deliberate” and produced the most change-of-mind decisions in the wait zone. Sweis et al. discovered behavioral evidence of compensatory valuations on subsequent trials compared to non-violation sequences much like Steiner and Redish, suggesting change-of-mind decision s following ballistic errors, too, can contribute to a post-decisional regret-like phenomenon.

Until now, it had been unexamined if these different operational definitions of regret published by Steiner and Redish 2014 (regret type I) versus Sweis et al. 2018 (regret type II) are related and share a common computational basis for generalized “mistake appraisal” or instead capture fundamentally distinct processes. Furthermore, it has not yet been tested if sensitivity to regret might be linked to individual differences in stress-response traits. Here, we directly compare both operational definitions of regret head-to-head in mice tested following one of two experimental manipulations: (i) challenging mice in a well-validated animal model used to study depression (chronic social defeat stress) capable of categorizing individuals into two classes of different stress-response phenotypes: stress-susceptible and stress-resilient; and (ii) virally knocking down the transcription factor CREB, a known regulator of stress-sensitive genes, in either the medial prefrontal cortex or nucleus accumbens – regions important for decision making and in which disrupting CREB function is known to promote stress-susceptible versus stress-resilient traits, respectively (*11, 15*). We report that neither social defeat stress nor viral CREB manipulations altered the ability to acquire the Restaurant Row task and had no effect on gross locomotor or feeding behavior. However, we found an increase in sensitivity to regret type I and a decrease to regret type II in stress-susceptible mice only. On the other hand, stress-resilient mice like non-defeated controls never displayed regret type I and instead displayed enhanced sensitivity to regret type II over that of non-defeated controls. Dissociable alterations in sensitivity to these types of regret could be induced by perturbing the function of CREB in the medial prefrontal cortex versus nucleus accumbens of stress-naïve mice. Taken together, we demonstrate that the effects of mistake history on influencing future decisions differentially depend on the nature of the erroneous choices made, these differences arise from dissociable brain structures, and can be extracted from mice with unique stress-response predispositions.

## Results

### Restaurant Row task

Here, C57BL/6J male mice were trained on the Restaurant Row task variant previously described in Sweis et al. 2018 (*7*). Mice had a limited daily time budget (60 min) to forage for their sole source of food by running counterclockwise around a square maze (Fig. 1a). Each corner “restaurant” was decorated with unique contextual cues and contained the reward site of a single, fixed flavor (chocolate, banana, grape, or plain). Flavors were used to modulate subjective value determined by revealed preferences without assuming reward value (as opposed to varying pellet number in each restaurant as this would come with added complexity due to increased handling time per pellet). Flavor preferences were determined by summing the end-of-session total pellets earned in each restaurant and ranking flavors from most preferred to least preferred (Fig. 1b). Upon entry into a restaurant’s T-shaped intersection (offer zone), an offer was randomly selected from a uniform distribution of 1 to 30 sec. The corresponding tone frequency associated with the selected delay sounded repeatedly until an explicit decision to enter the wait zone was made. If mice chose to enter the wait zone, tones descended stepwise with decreasing pitch counting down progress toward obtaining the reward. Alternatively in the offer zone, mice could choose to skip the wait zone and instead advance down the hallway to the next restaurant, as mice were required to encounter restaurants serially.

**Figure 1.**
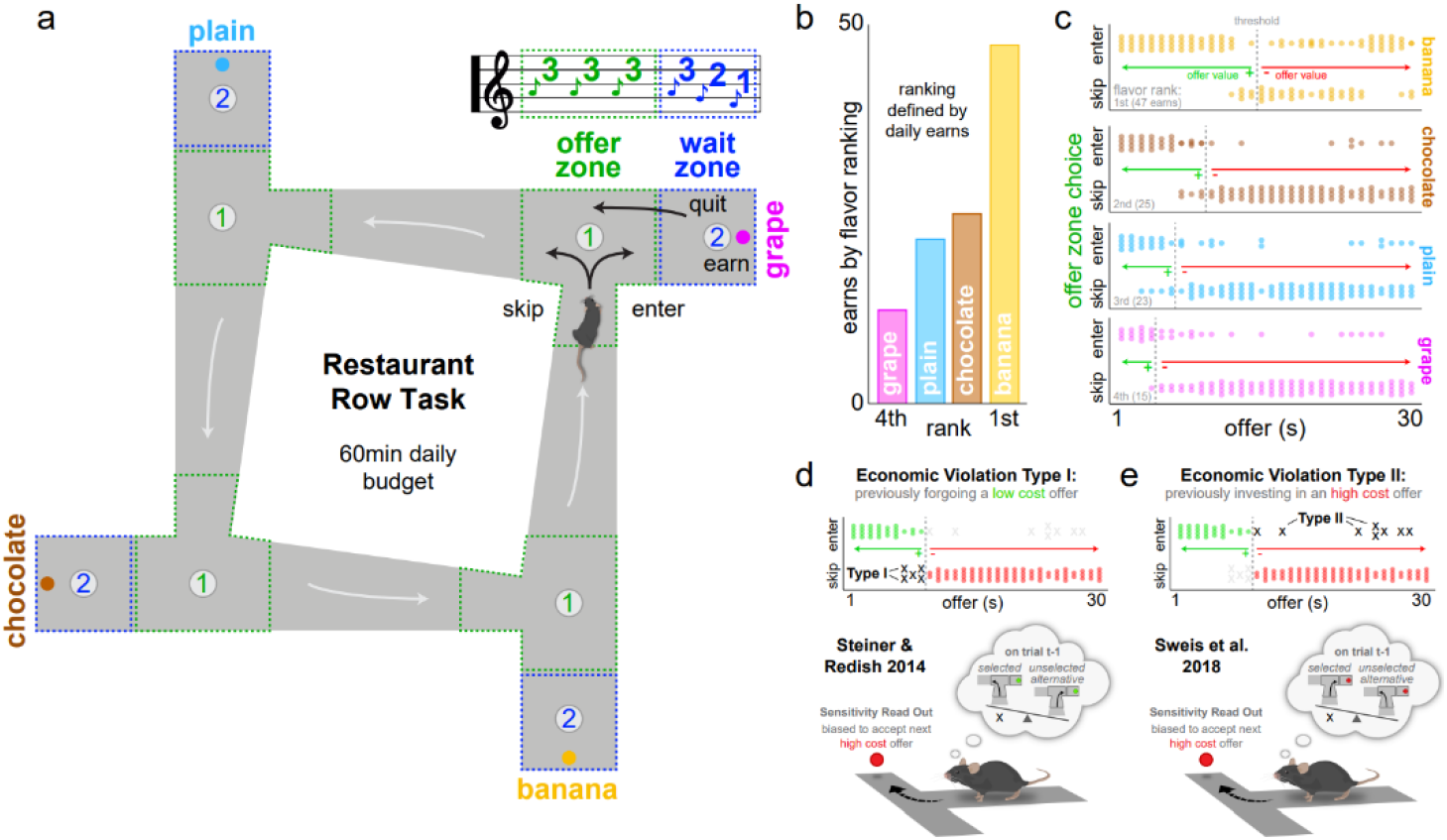
Restaurant Row task. (a) Task schematic. Mice had 60 min to forage for their only food for the day by making serial decisions in each uniquely flavored restaurant. Reward delays were randomly selected from a range of 1 to 30 s and signaled by the pitch of a tone. Trials ended if mice skipped in the offer zone, quit during the tone countdown in the wait zone, or earned a pellet before being required to advance to the next restaurant. (b) Pellets earned in each restaurant determine daily flavor rankings. Example session from a single mouse. (c) Economic thresholds of willingness to wait were calculated by fitting a logistic regression to whether or not food was earned as a function of cued offer cost and identifying the inflection point of each curve. Example session from a single mouse. Dots represent individual trials. Vertical dashed lines indicate thresholds in each restaurant. Thresholds determine offer value [V_offer_ = threshold – offer]: offers below threshold reflect positively valued offers (green arrow) that are typically earned while offers above threshold are negatively valued (red arrow) and typically skipped. (d-e) Two types of economic violations in the offer zone are defined by atypical decisions to skip a low cost, high value offer (d, type I) or enter a high cost, low value offer (e, type II). Both types of violations have been previously demonstrated to bias animals to make compensatory decisions on subsequent trials relative to non-violation decisions – a behavioral readout of sensitivity to regret-related processes.

Threshold s of willingness to wait were calculated in each restaurant by fitting whether or not mice earned rewards to a sigmoid as a function of cued offer cost and identifying the inflection point of each curve (Fig. 1c). This metric reflects the cost of an offer below which mice typically earn and above which mice typically forgo. Thus, the value of an offer on a given trial can be calculated by subtracting the delay from one’s threshold derived from that day’s session. This allows offers across mice with individual differences in subjective flavor preferences and across days to be normalized to one’s own indifference point. Economic violations on this task were defined here as atypical choices that violate one’s own stable decision policies. Skipping an offer below threshold (offer value > 0) constitute what we define here as “economic violation type I” and capture the events investigated in Steiner and Redish 2014 (Fig. 1d) (*5*). Accepting an offer above threshold (offer value < 0) constitute what we define here as “economic violation type II” and capture the events investigated by Sweis et al 2018 (Fig. 1e) (*7*). Because time is a limited commodity on this task, decisions are interdependent across trials. Thus, sensitivity to the consequences of having made these distinct types of economic violations were analyzed on the subsequent trial using the behavioral read out previously published: bias to accept a high-cost offer on the subsequent trial (Fig. 1d-e) (*5, 7*). This metric captures how mistake history can bias individuals to subsequently make atypical or compensatory reward-seeking decisions. We examined these behaviors in mice following one of two experimental manipulations: (i) following chronic social defeat stress or (ii) following brain-region-specific molecular manipulations.

### Chronic social defeat stress manipulation

In the first cohort of mice, we subjected animals to the chronic social defeat stress protocol that effectively distinguishes between stress-susceptible (SUS) and stress-resilient (RES) animals alongside non-stressed controls (CON, Fig. 2a) (*15*). C57BL/6J mice were exposed to aggressive CD-1 male mice daily for 10 consecutive days. Animals were allowed to physically interact for approximately 5 min before being separated by a mesh partition. They the n remained cohoused for the rest of the day to allow for continuous sensory interaction before repeating defeat with a different CD-1 mouse. Following the final defeat, C57BL/6J mice were tested in a rapid social interaction screening assay measuring approach versus avoid behavior toward a novel CD-1 mouse. Social avoidance induced by this protocol has served as a well-validated predictor of several additional depression-related phenotypes on numerous other rapid behavioral screening tests and thus is commonly used as a metric to define SUS mice (Fig. 2b) (*15*). CON mice on the other hand typically approach the novel CD-1 mouse during the social interaction screen. Therefore, RES mice are commonly defined as those individuals who, too, engage in approach behavior similar to CON mice.

**Figure 2.**
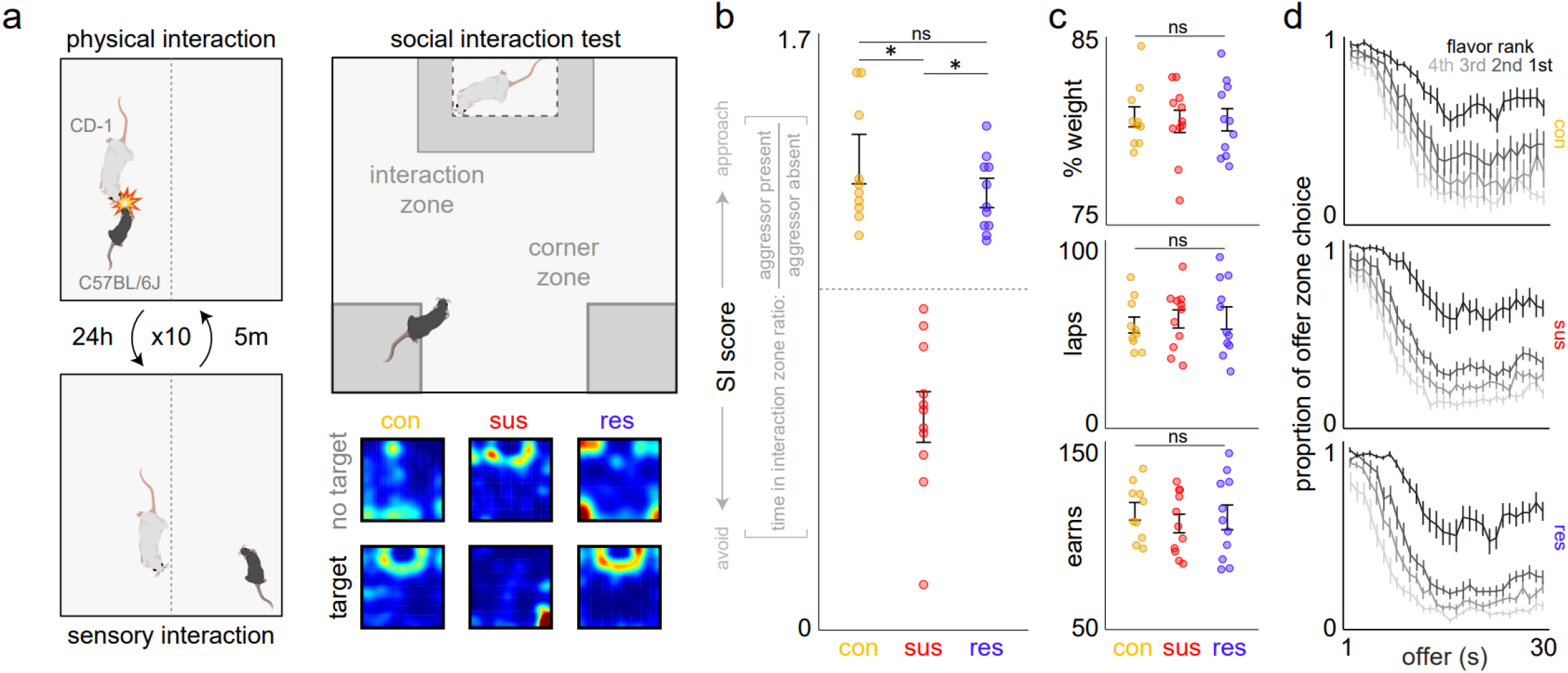
Chronic social defeat stress. (a) Stress protocol. C57BL/6J mice were exposed to chronic social defeat stress by being paired with aggressive CD-1 male mice, while non-stressed controls (CON, n=10) were instead paired with each other. After 5-min daily physical interactions followed by constant visual, olfactory, and auditory contact for 10 days with different CD-1 mice, C57BL/6J mice were tested on a social interaction (SI) assay. Heat maps show time in the interaction zone (max 3 min) from example sessions with and without a CD-1 target mouse present. (b) SI score calculated from the ratio of time in interaction zone with CD-1 target present to when the target was absent, capturing approach versus avoidance behavior. SI scores above and below 1 classified defeated mice into stress-resilient (RES, n=11) and stress-susceptible (SUS, n=11) groups, respectively (one-way ANOVA: *F*_2,31_=47.975, \**p*<0.0001, Tukey_CON/RES_ *t*=1.15, ^ns^*p*=0.495). (c) No differences between groups in average pre-task percentage baseline body weight (top, ANOVA: *F*_2,31_=0.117, ^ns^*p*=0.89), average number of daily laps run on the task (middle, *F*_2,31_=1.371, ^ns^*p*=0.27), and average total number of daily pellets earned on the task (bottom, *F*_2,31_=0.515, ^ns^*p*=0.60). (d) Probability of enter decisions made while in the offer zone as a function of cued offer cost split by restaurant ranked by subjective flavor preferences. See Fig. S1a for quantification of thresholds, or indifference points, by restaurant. Dots represent individual animals. Error bars ± 1 SEM. Not significant (ns).

Following this protocol, all mice were food restricted for 3 days to approximately 80% body weight and trained on the Restaurant Row task. All mice acquired the basic structure of the task. Mice were well trained until their performance stabilized across several behavioral metrics, including body weight, laps run, and pellets earned. Defeat had no immediately observable effects on overt locomotor or feeding behavior (Fig. 2c). SUS, RES, and CON mice ran equivalent number of laps and earned the same amount of food remaining stable at similar body weights. All mice learned to reliably discriminate tones by accepting low cost offers and rejecting high cost offers (Fig. 2d). Importantly, because mice also treated the same tones differently in each restaurant indicates that animals understood the economic structure of the task. That is, mice made informed economical decisions that integrate delay information communicated via auditory cues with flavor information communicated by visuospatial cues to forage effectively while also prioritizing subjective preferences. Mice readily revealed subjective flavor preferences whose ordinal rankings among the four flavors as indicated by thresholds of willingness to wait were matched across SUS, RES, and CON mice (Fig. S1a).

### Distinct types of economic violations impact future choices

Next, we examined how each type of economic violation influenced subsequent decisions on this task. Economic violation type I is defined by situations in which mice skip a positively valued offer (offer below one’s threshold). Importantly, regret could be induced following this choice if mice subsequently encounter a negatively valued offer (offer above one’s threshold) on the next trial (Fig. 3a). This critical distinction separates a mere mistake or atypical choice from one that may be specifically linked to a regret-related process. Thus, this sequence operationalizes a risky decision in a foraging task that results in a poor outcome and is highlighted by a missed opportunity when information is provided on the second trial. To simplify the labeling of these trials, we will refer to the first trial, the trial on which the violation occurred, as “trial t-1.” In order to quantify how mistake history on trial t-1 might bias mice to subsequently make atypical decisions, we calculated the probability of accepting the negatively offer on the second trial, offers animals would typically reject. We call this trial, the trial following the violation, the “read-out trial.” These specific sequences were not constructed a priori nor built into the task design but rather identified post-hoc extracted from an animal’s natural encounters while foraging among random offers that were uniformly distributed. The bias toward or probability of accepting offers on the read-out trial is determined by dividing the number of enter choices on this trial by total number of offers presented on this trial given that (i) the offer on trial t-1 was positively valued, (ii) the choice on trial t-1 was a skip decision, and (iii) the offer on the read-out trial was negatively valued (see Fig. S2 for visual explanation). In order to control for error of one’s own agency – a critical tenant of regret – enter bias on the read-out trial following this violation sequence was compared to a matched economic scenario in which animals did not commit a violation of their own decision policy on trial t-1. This control sequence therefore was defined by the same offers (positive offer value on trial t-1 and negative offer value on the read-out trial) but only differed in that an economically congruent enter decision was made on trial t-1. A difference in decision bias on the read-out trial between the violation and control sequences allows us to behaviorally measure sensitivity to a choice-history-dependent regret-related process. Because these scenarios depend on sequences between two restaurants, we split this analysis by each restaurant aligned to the ranking of the restaurant on the read-out trial. This disambiguates the different enter rates across restaurants on the read-out trial that go into this analysis and allows for a closer examination of how relative subjective value between restaurants interacts with choice history, which has been previously shown on this task to anchor one’s decision bias (*7*). By comparing these two sequences, we found that mice displayed an increase in the probability of entering negatively valued offers on the read-out trial following economic violation type I compared to non-violation control decisions on trial t-1 (Fig. 3b, Fig. S3a). These data reproduce findings from Steiner and Redish 2014 and indicate that the poor outcome of a risky skip decision can drive an individual to be more likely to make a subsequent choice they typically would not (*5*). Surprisingly, in the present study we found that this is a property unique to SUS mice but not CON or RES mice (Fig. 3b, Fig. S3a). Importantly, the ability to detect this effect required fully leveraging the economic complexity of Restaurant Row separating the value space across the three primary dimensions of the task: offer value, restaurant identity, and choice history. Furthermore, violation rates on trial t-1 were not different between restaurants or groups of mice (Fig. S4a), indicating behavioral differences were not due to frequency of scenarios encountered or willingness to “underspend” on trial t-1, but rather how animals weigh choice history differently during subsequent decisions.

**Figure 3.**
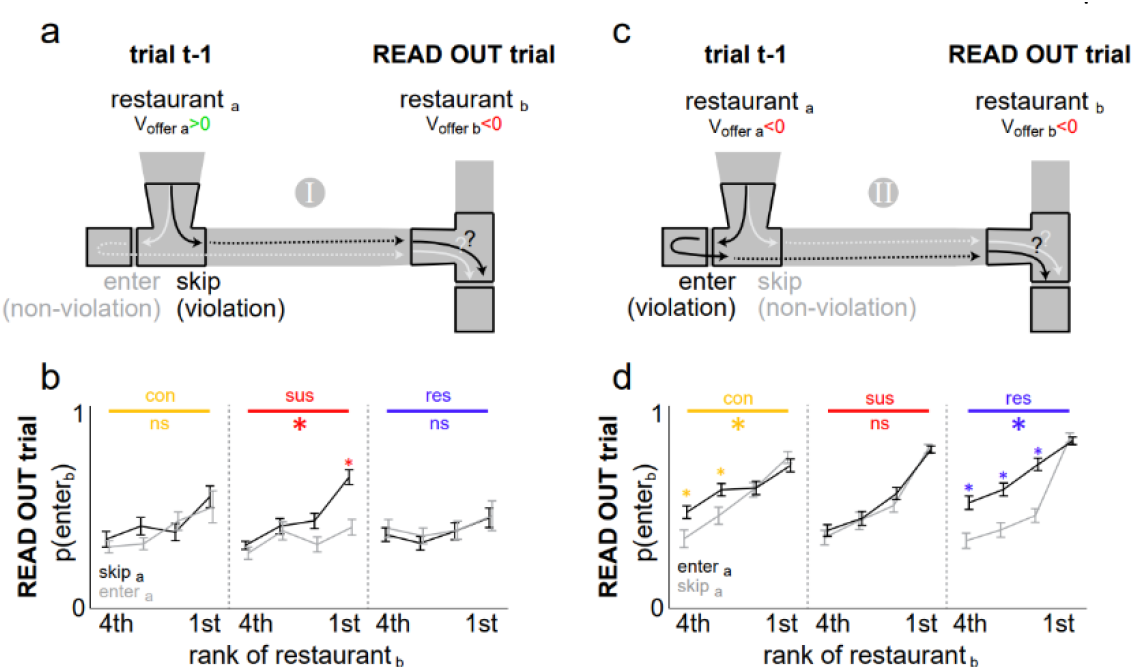
SUS and RES mice display differential changes in behavior following two distinct types of economic violations. (a) Sequence schematic involving economic violation type I on trial t-1 (skip positively valued offer, black arrow) followed by a negatively valued offer presented on the subsequent read-out trial. Control sequence varies only in that a non-violation decision was made on trial t-1 (enter positively valued offer, gray arrow). (b) The probability of entering the offer presented on the read-out trial is plotted in both the violation and non-violation sequences, split by restaurant aligned to the read-out trial and ranked by subjective flavor preferences. See Fig. S2 for a visual explanation of this analysis. (c) Sequence schematic involving economic violation type II on trial t-1 (enter negatively valued offer, black arrow) followed by a negatively valued offer presented on the subsequent read-out trial. Control sequence varies only in that a non-violation decision was made on trial t-1 (skip negatively valued offer, gray arrow). (d) Decision bias on the read-out trial as described in (b) for economic violation type II and non-violation control sequences. See Fig. S3a-b for quantification of the difference score in the decision bias on the read-out trial between violation and non-violation sequences. Error bars ± 1 SEM. Not significant (ns).

Economic violation type II is defined by situations in which mice enter a negatively valued offer (offer above one’s threshold) on trial t-1. Similar to the above analysis, we calculated the probability of accepting a negatively valued offer on the read-out trial following this type of violation (Fig. 3c). Thus, the bias toward or probability of accepting offers on the read-out trial is determined by dividing the number of enter choices on this trial by total number of offers presented on this trial given that (i) the offer on trial t-1 was negatively valued, (ii) the choice on trial t-1 was an enter decision, and (iii) the offer on the read-out trial was negatively valued. Here, the non-violation decision on trial t-1 would have been to skip such trials instead and thus serves as the control sequence to compare how choice history in this economic scenario can influence subsequent decisions. We found that mice displayed an increase in the probability of entering negatively valued offers on the read-out trial following economic violation type II compared to non-violation control decisions on trial t-1 (Fig. 3d, Fig. S3b). These data reproduce findings from Sweis et al. 2018 and indicate that erroneous decisions to accept economically disadvantageous offers, too, can drive an individual to be more likely to make a subsequent choice they typically would not (*7*). Interestingly, while this effect was observed in CON mice to a similar degree consistent with previous reports (*7*), it was significantly enhanced in RES mice and surprisingly completely absent in SUS mice (Fig. 3d, Fig. S3b). This effect, too, was unrelated to scenario frequency, as violation type II rates on trial t-1 were not different between restaurants or groups of mice (Fig. S4b). Furthermore, this pattern of effects was not seen following violations on trial t-1 if a positively valued offer was instead presented on the read-out trial (Fig. S5a-b), suggesting the ability to detect these mistake-related phenomena depend on subsequently probing animals with an economically disadvantageous offer.

### Behavioral analyses of choice processes

In order to better understand the processes involved in these economic violations, we examined how mice executed decisions in the offer zone on trial t-1. Because path trajectories can reveal underlying decision-making processes, we analyzed body positions during passes through the offer zone choice point leading up to the decisions on trial t-1. Ballistic versus tortuous trajectories can serve as behavioral proxies of different neurophysiological processes governing these choices. Vicarious trial and error (VTE) behavior captures pause-and-look reorientation events that correlate with alternating neural representations of competing paths forward ahead of the animal, thought to be part of a prospective planning or deliberation process (*17-19*). VTE measures the absolute integrated angular velocity of a pass through a choice point. The amount of physical “hemming and hawing” is best quantified by computing changes in velocity of every *x* and *y* body position grabbed from each video frame stepped over time as *dx* and *dy*. From these vectors, we calculated the instantaneous change in angle, *Phi*, as *dPhi* that is then integrated over the pass through the offer zone from offer onset until either a skip or enter decision was made as *IdPhi*.

We found that VTE was significantly higher for skip decisions than enter decisions (Fig. 4a-b), consistent with previous report suggesting offer-skipping behaviors on this task invoke more hesitation (*7, 20*). Conversely, enter decisions instead are generally low VTE events that are typified by ballistic journeys through the choice point. Based on the heading direction of skip paths, skip decisions chiefly comprise near-enter trajectories that are re-routed mid-journey in the offer zone, suggesting VTE incorporates a delayed valuation to override prepotent or habit-like offer-taking responses on this task. As a function of offer value, VTE generally displays an inverted U-shaped curve with a left-shifted peak such that negatively valued offers just above one’s thresholds elicit the most VTE (Fig. 4c). This suggests that the most difficult to process decisions are for offers that are just more expensive than one is willing to wait, consistent with previous reports (*7, 20*). Interestingly, the probability of appropriately skipping negatively valued offers increases as a function of the amount of VTE displayed in the offer zone (Fig. 4d). This indicates that the outcome of such a deliberative process is more likely to result in an economically advantageous decision the more an animal engages in VTE. Furthermore, the amount of VTE required to reliably skip negatively valued offers is higher in more preferred restaurants (Fig. 4d). Taken together, these data indicate that choices in the offer zone can employ a flexible decision-making process that integrates conflict between expensive although desirable rewards. These data also indicate that a failure to engage in VTE – what we term “snap-judgments” – can contribute to type II violations, unlike type I violations that follow from high VTE events. Surprisingly, SUS mice did not display an inverted U-shaped VTE curve and instead showed higher VTE when encountering positively valued offers compared to RES and CON mice (Fig. 4c). Additionally, RES mice displayed less VTE when encountering negatively valued offers and required less VTE to skip negatively valued offers compared to SUS and CON mice (Fig. 4c-d). These findings suggest that SUS and RES mice may be attending to or integrating information throughout their offer zone decision process differently depending on the value of the offer, which could differentially impact the weight each type of economic violation carries into future choices.

**Figure 4.**
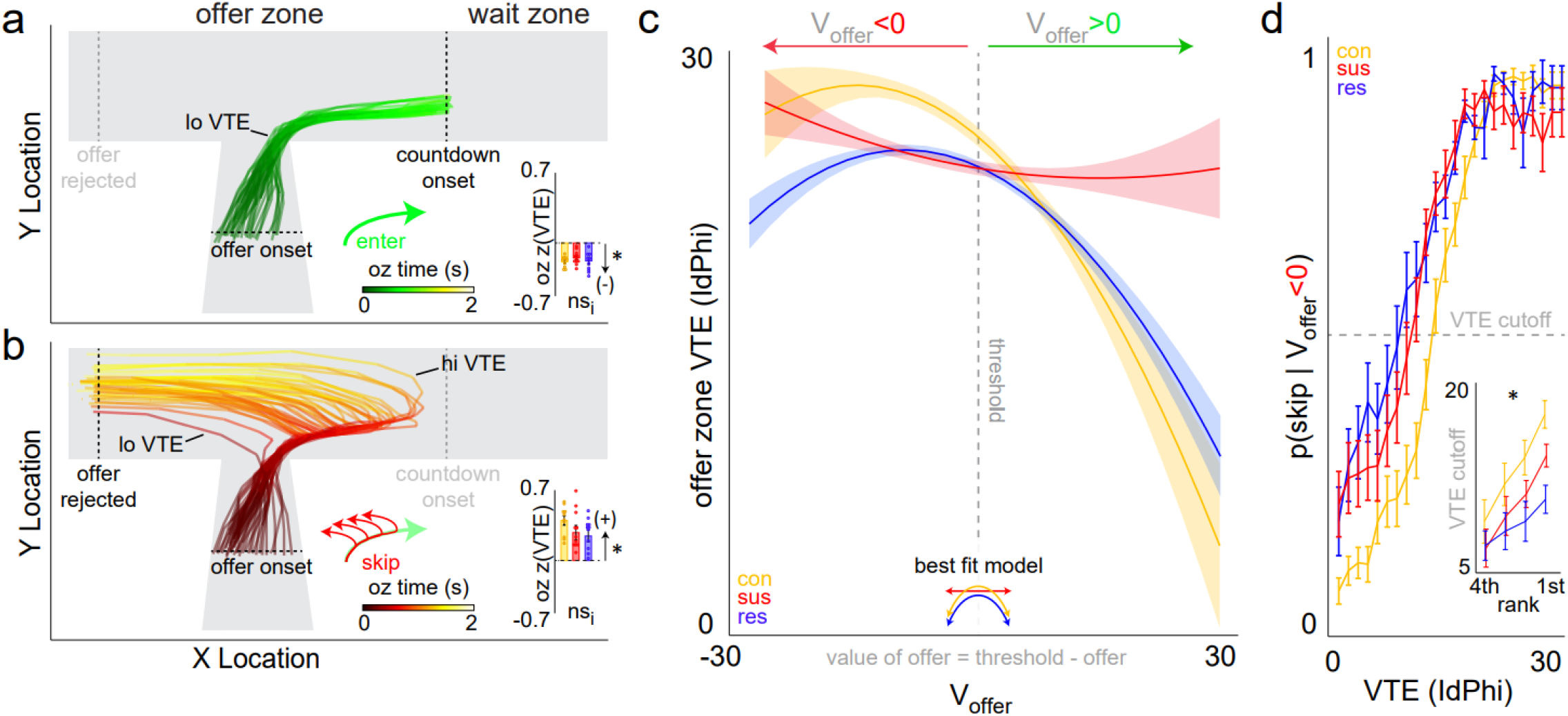
SUS and RES mice display value-based differences in vicarious trial and error (VTE) behavior in the offer zone. (a-b) Traces of overlayed path trajectories in a single restaurant taken from an example session separated by offer zone (oz) decisions to (a) enter or (b) skip. Trajectory traces through the offer zone is color-scaled using a heat map that represents time from offer onset until either an enter or skip decision was made. Examples of lo versus hi VTE trials are labeled. Note: skip paths often involve re-routing of near enter trajectories. Insets display z-scored VTE behavior across groups of mice (CON - yellow, SUS - red, and RES – blue; sign test, enter: *t*_31_=-6.883, \**p*<0.0001; skip: *t*_31_=+8.431, \**p*<0.0001; one-way ANOVA, enter: *F*_2,31_=0.131, ^nsi^*p*=0.878; skip: *F*_2,31_=1.684, ^nsi^*p*=0.203). (c) VTE behavior in the offer zone as a function of offer value [V_offer_ = threshold – offer]. Vertical dashed line indicate threshold of willingness to wait. Icon inset displays the model of best fist when comparing linear versus quadratic functions. AIC Weights: CON – ^ns^linear: 1.567×10^−14^, *quadratic: 0.999; SUS – *linear: 0.718, ^ns^quadratic: 0.282; RES – ^ns^linear: 4.81×10^−4^, *quadratic: 0.999. (d) Probability of skipping negatively valued offers in the offer zone (i.e., avoiding type II violations) as a function of VTE behavior. Horizontal dashed line indicates the amount of VTE required to skip at least 50% of such trials (VTE cutoff). Inset displays average VTE cutoff split by ranked flavor preferences. Significant interaction between groups and rank (two-way ANOVA: *F*_2,3_=2.837, \**p*<0.05). Error bars ± 1 SEM. Not significant (ns).

Next, we examined how mice executed decisions in the wait zone on trial t-1 in order to better understand the behavioral sequala following economic violation type II: accepting negatively valued offers. In the wait zone, mice were tasked with remaining near the feeding site during the tone countdown after making an enter decision in order to earn a reward but were free to quit at any moment. We found that the vast majority of quits occurred following enter decisions for negatively valued offers (Fig. 5a). Furthermore, we found that the point at which mice quit negatively valued offers occurred most frequently with an amount of time remaining in the countdown that was above one’s threshold (Fig. 5a). That is, the value of the time left required to obtain the reward of a negatively valued offer, too, was likely still negative at the time of quitting. This indicates that many type II violations in the offer zone resulted in wait zone quit decisions and that the majority of these quit decisions in the wait zone were economically advantageous choices that effectively corrected offer zone violations – decisions that were the product of a failure to engage in VTE. This allows us to confidently label enter decisions for negatively valued offers indeed as “mistakes.” Consistent with previous reports, these data highlight how re-evaluating recent mistakes and change-of-mind decisions following ballistic events can contribute to post-decisional regret (*17-19, 21*). SUS, RES, and CON mice all executed quit decisions in this manner and to similar degrees (Fig. 5b), suggesting not overall quit frequency but rather how animals weigh the value of choice history following these violations is likely altered. Previous reports have shown that a closer examination of the quitting process can reveal hidden costs during change-of-mind decisions and could shed light on how animals may be differently valuing future reward-seeking behavior following type II violations (*22*).

**Figure 5.**
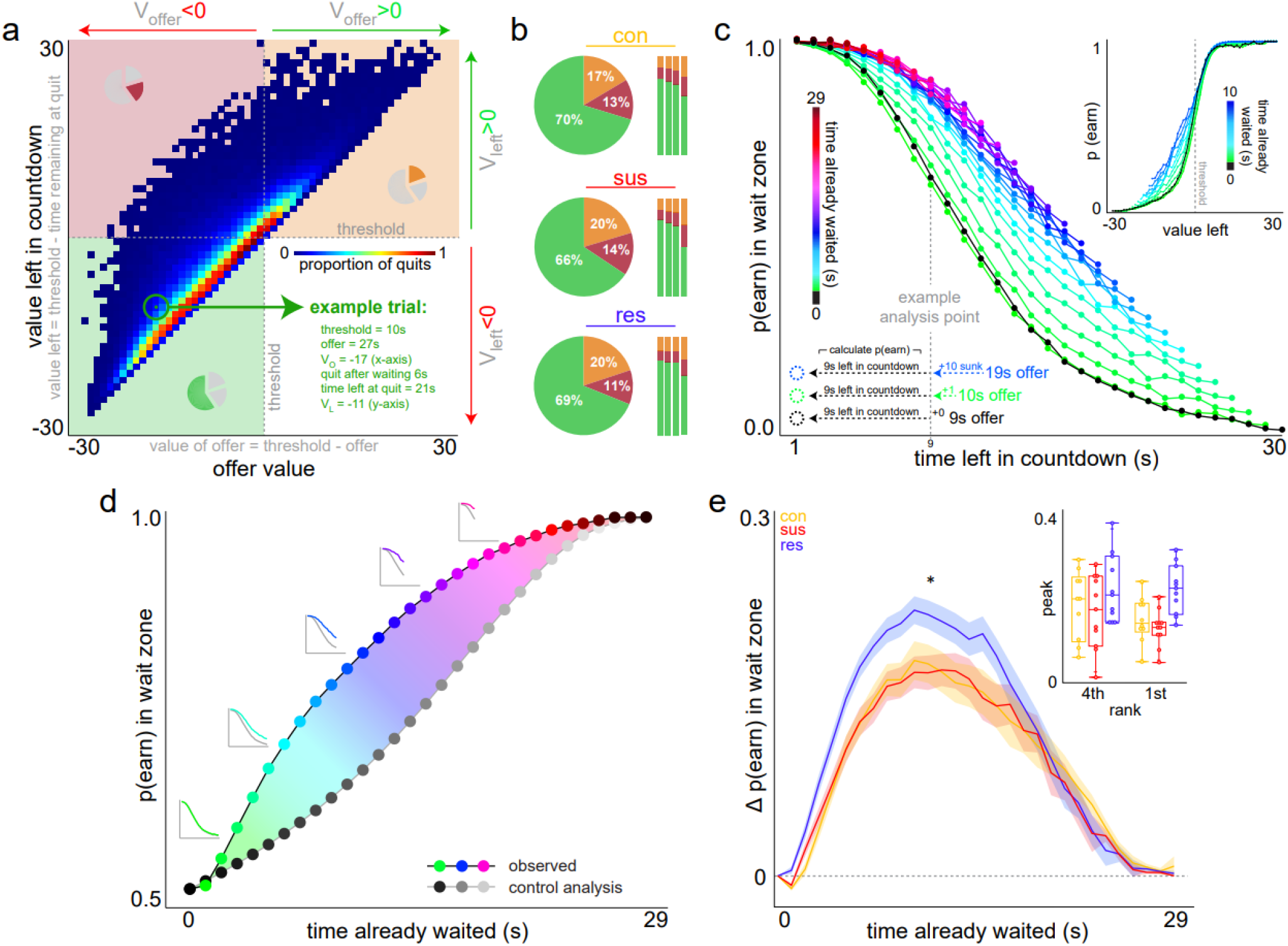
RES mice display enhanced sensitivity to sunk costs during change-of-mind quit decisions in the wait zone. (a) Quit decisions in the wait zone separated by offer value [V_offer_ = threshold – offer] and the value left in the countdown at the moment of quitting [V_left_ = threshold – time remaining]. Note: trials where V_offer_>0 & V_left_<0 do not exist). Heatmap of quit trials shows the full distribution of change-of-mind decisions across these value dimensions. Each pixel represents a value combination pooled across all animals. Pie inserts show the occupancy of each value quadrant relative to all quits. Note: the most common type of quit decision is for V_offer_<0 & V_left_<0 (green quadrant). Pixel circled in green describes an example trial. (b) Relative proportion of quit types across the three quadrant categories defined in (a) split by groups of mice. All quit trials (left, pie) and relative to each restaurant (right, bars) ranked from least preferred to most preferred flavors left to right. (c) Sunk cost analysis. Probability of earning in the wait zone as a function of time left in the countdown (x-axis) and time already waited (color). Example analysis text overlay describes three example sunk cost conditions: 0 s (black), 1 s (green), and 10 s (blue) all matched for the same amount of time left in the countdown. Thus, each sunk cost condition arose from trials with different starting offers (9, 10, and 19 s, respectively). Probability of earning was calculated in these three examples based on the number of trials remaining at the 9 s left time point but using all combinations of time spent - time left permutations (See Fig. S6a) for the full spectrum analysis. Data pooled across all animals. Inset displays the analysis aligned to threshold (vertical dashed gray line) as a function of V_left_ for the first 10 s of time already waited. Note: the sunk cost effect is present only for V_left_<0. (d) By collapsing across the time left axis in (c), the dimensions of this analysis can be simplified, capturing the envelope of the overall effect of time already waited, or sunk costs, on p(earn). Each colored curve in (c) is averaged into a single datapoint in (d), with 5 example insets replotted for visualization of 1, 5, 10, 15, and 20 s sunk cost conditions. To control for missing data points when reducing dimensions at each sunk cost condition, data points from the 0 s sunk cost condition in (c, black curve) were reiteratively collapsed upon itself leaving out the right most value at each step to match the number of datapoints and starting offers from the observed sunk cost conditions (inset, gray traces). Data pooled across all animals. (e) The difference between the two resulting curves in (d) captures the overall magnitude of sensitivity to sunk costs. Horizontal dashed gray line indicates zero difference score between curves in (d). Two-way ANOVA, main-effect_sunk_: *F*_29,31_=69.992, \**p*<0.0001, interaction_groupXsunk_: *F*_2,29_=11.614, *^i^*p*<0.0001. Error bars ± 1 SEM. Not significant (ns).

Regret derived from economic violation type II – atypically accepting a high-cost offer and being faced with the dilemma to change one’s mind during continued investment – is related to a well-studied human cognitive bias known as the sunk cost fallacy (*22*). This describes the phenomenon during which irrecoverable losses can escalate the commitment of an ongoing endeavor, even if suboptimal, and is thought to generate cognitive dissonance on some level (*22-24*). In the present task, quit decisions capture the continuous re-evaluation of an ongoing investment paid toward earning a reward. Following enter decisions for negatively valued offers, spending time to make an efficient quit decision competes in a race toward the threshold after which point it would be optimal to finish waiting as mice anticipate an approaching goal. Time already spent, or sunk costs, when deciding to quit therefore is a rolling economic entity that should not but is indeed capable of carrying compounding value over the passage of time that opposes quitting and can pressure an animal to continue waiting. We developed a dynamic analysis capable of extracting the hidden effects of sunk costs on the likelihood of quitting that has been previously published in rodents and humans tested on translated versions of the Restaurant Row task (*22*). This analysis separates the time already waited during a given countdown in the wait zone from future time left required to obtain a reward on that trial and measures how these two dimensions of time independently accumulate value that promotes staying in the wait zone. Each quit trial was parsed into bins of [time spent, time left] pairs at the moment of quitting from which many permutations arise based on various starting offers (Fig. S6a). The probability of earning a reward in the wait zone was dynamically calculated along a continuum using a sliding window survival analysis as both a function of time left in the countdown as well as time already waited (Fig. 5c).

Consistent with previous reports across species, we found that the time already waited increases the likelihood of continuing to wait to earn a reward independent of the temporal distance to the goal (Fig. 5c-e) (*22*). Additionally, the more time that was already waited, the stronger this effect, a critical tenant of the sunk cost phenomenon (Fig. 5c-e). Interestingly, this effect is largely driven by time spent waiting after accepting negatively valued offers compared to equivalent time spent waiting after accepting positively valued offers (Fig. 5c). While all mice were sensitive to the effects of time already waited on the value of staying, this phenomenon was uniquely robust in RES compared to SUS or CON mice (Fig. 5e). Neither the amount of time spent in a restaurant’s offer zone before making type II violations nor time spent since the last reward was earned influenced the likelihood of waiting once in the wait zone for all mice (Fig. S6b-d). These data indicate that time spent considering quitting negatively valued offers carries unique weight that is enhanced in RES mice. Taken together, these data suggest that how type I and type II violations influence future decisions for SUS, RES, and CON mice may be linked to valuation differences in the decision-making processes of the mistakes themselves.

### Brain-region-specific CREB manipulation

Having demonstrated that different operational definitions of regret as described by Steiner and Redish 2014 and Sweis et al. 2018 do not always covary but rather are indeed separable following a behavioral manipulation, in the second cohort of mice we aimed to perturb these processes with a biological manipulation. Here, we probe two brain structures important for value-based decision making and engaged by the Restaurant Row task in both rodents and humans. Specifically, we targeted the medial prefrontal cortex (mPFC) and nucleus accumbens (NAc) as each region has been shown to be involved in processes related to type I and type II economic violations (*5, 25-27*). To date, there are no studies linking molecular neuroscience at the level of gene regulation to neuroeconomics.

Here, we inhibited CREB function in mPFC or in NAc neurons via viral-mediated overexpression of a dominant negative CREB mutant (mCREB) in stress-naïve mice, and then trained these animals in Restaurant Row (Fig. 6a, Fig. S8). Control animals were transfected with a GFP-only virus in either region. Following these surgeries, mice were allowed to recover and then were food restricted for 3 days to approximately 80% body weight before being trained on the Restaurant Row task. Well-trained mice ran equivalent number of laps and earned the same amount of food remaining stable at similar body weights (Fig. 6b). All mice were capable of reliably discriminating tones as a function of cued offer cost and revealed subjective flavor preferences whose ordinal rankings among the four restaurants as indicated by thresholds of willingness to wait were matched across groups (Fig. 6c, Fig. S1b). We found that, compared to GFP controls, expression of mCREB in the mPFC or NAc increased sensitivity to type I economic violations (Fig. 6d-e, Fig. S3c): following skip decisions for positively valued offers (violation) compared to enter decisions for similar offers (non-violation) on trial t-1, these animals displayed an increased likelihood of accepting negatively valued offers on the read-out trial. These data suggest normal CREB function in either region is required to suppress regret-related processes associated with this type of economic violation. Interestingly, compared to GFP controls, we also found that expression of mCREB only in the mPFC increased sensitivity to type II economic violations (Fig. 6f-g, Fig. S3d): following enter decisions for negatively valued offers (violation) compared to skip decisions for similar offers (non-violation) on trial t-1, these animals displayed an increased likelihood of accepting negatively valued offers on the read-out trial. In contrast, expression of mCREB in the NAc decreased sensitivity to type II economic violations. Groups did not differ in their frequency of type I or type II violations (Fig. S4c-d) or if a positively valued offer was instead presented on the read-out trial (Fig. S5e-h). Importantly, these data reveal that sensitivity to different types of economic violations are capable of being modulated independently depending on the brain region perturbed and rather than share a generalized basis for “mistake appraisal” instead capture separable, fundamentally distinct action-specific computational processes.

**Figure 6.**
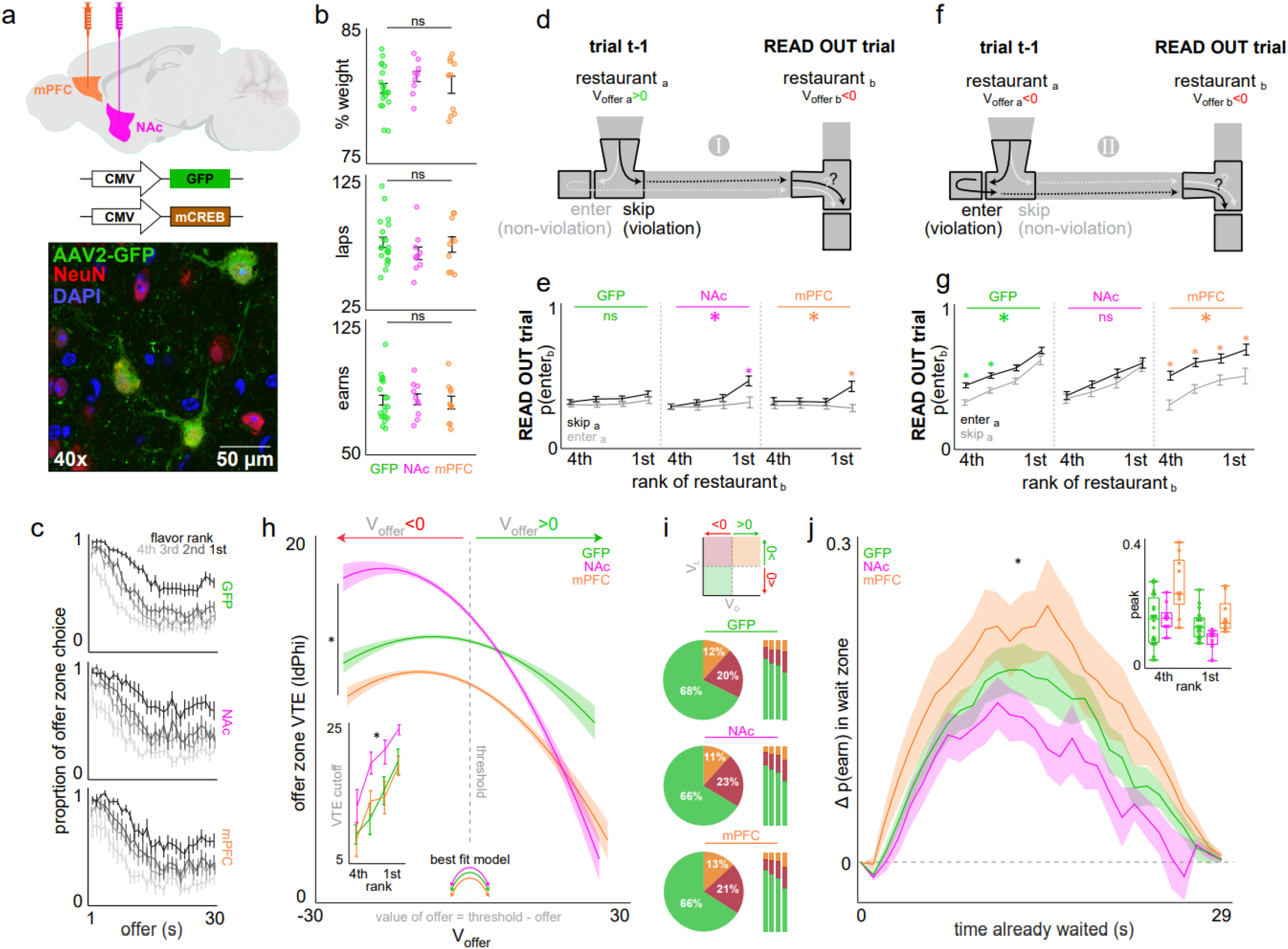
CREB function in the mPFC and NAc differentially influences sensitivity to distinct types of economic violations with associated changes in offer zone VTE and wait zone sunk cost behavior. (a) Mice were transfected in either the mPFC (n=10) or NAc (n=10) with a neurotropic adeno-associated virus (AAV2) encoding either green fluorescent protein (GFP, n=20) or a dominant-negative mutant form of CREB (CREBS133A, mCREB). Representative confocal image taken from sectioned NAc tissue (bottom). See Fig. S7 for viral targeting. (b) Average pre-task percentage baseline body weight (top, ANOVA: *F*_2,31_=0.033, ^ns^*p*=0.97), average number of daily laps run on the task (middle, *F*_2,31_=0.718, ^ns^*p*=0.49), and average total number of daily pellets earned on the task (bottom, *F*_2,31_=0.190, ^ns^*p*=0.83). (c) Probability of enter decisions made while in the offer zone as a function of cued offer cost split by restaurant ranked by subjective flavor preferences. See Fig. S1b for quantification of thresholds, or indifference points, by restaurant. (d-g) sequence schematics (d, f) and decision bias on the read-out trial (e, g) as described in Fig. 3 for economic violations type I (d, e) and II (f, g) with respective non-violation control sequences. See Fig. S2 for a visual explanation of this analysis. See Fig. S3c-d for quantification of the difference score in the decision bias on the read-out trial between violation and non-violation sequences. (h) VTE behavior as a function of offer value [V_offer_ = threshold – offer]. Vertical dashed line indicate threshold of willingness to wait. All groups are better explained by a quadratic (displayed icon) than a linear function (NAc – ^ns^linear: 2.203×10^−23^, *quadratic: 0.999; mPFC – ^ns^linear: 5.318×10^−17^, *quadratic: 0.999; ^ns^GFP – linear: 0.083, *quadratic: 0.917), but NAc animals display more VTE (*F*_2,39_=123.443, \**p*<0.0001; Tukey_NAc/GFP_ *t*=+6.50, \**p*<0.0001) while mPFC animals display less VTE (Tukey_mPFC/GFP_ *t*=-15.18, \**p*<0.0001) on V_offer_<0 trials compared to GFP animals. This is partly reflected in increase for NAc mice ain the amount of VTE required to skip at least 50% of trials where V_offer_<0 (VTE cutoff, inset) split by ranked flavor preferences (significant main effect between groups *F*_2,39_=12.359, \**p*<0.0001; Tukey_NAc/GFP_ *t*=+3.28, \**p*<0.01; Tukey_mPFC/GFP_ *t*=-0.04, ^ns^*p*=0.999). (i) Relative proportion of quit types across the three categories represented by icon (top) defined in Fig. 5a split by groups of mice. All quit trials (left, pie) versus relative to each restaurant (right, bars) ranked from least preferred to most preferred flavors from left to right. (j) Sensitivity to sunk costs as described in Fig. 5e (*F*_2,29_=29.991, \**p*<0.0001). Dots represent individual animals. Error bars ± 1 SEM. Not significant (ns).

When examining how mice executed decisions on trial t-1, we found that mCREB expression caused several changes in both offer zone and wait zone behaviors. In the offer zone, all mice displayed an inverted U-shaped curve of VTE behavior as a function of offer value (Fig. 6h). However, compared to GFP controls, mCREB expression in either the mPFC or NAc decreased the amount of VTE mice displayed for positively valued offers. Interestingly, mCREB expression bidirectionally altered VTE for negatively valued offers in the mPFC versus NAc. mCREB expression in the mPFC decreased VTE for negatively valued offers while mCREB expression in the NAc increased VTE. Additionally, NAc-treated animals required a higher amount of VTE to appropriately skip negatively valued offers. Overall, the brain-region-specific effects of mCREB expression on VTE resulted in asymmetric changes in the offer zone depending on the value of the offer presented (i.e., shared direction of change for positively valued offers versus opposing direction of change for negatively valued offers). In the wait zone, all mice engaged in change-of-mind decisions most frequently following enter choices for negatively valued offers and when the value of the amount of time left in the countdown was still negative (Fig. 6i). Additionally, all mice were sensitive to how the amount of time already spent waiting for a reward, or sunk costs, increased the likelihood of staying in the wait zone independent of temporal distance to the goal (Fig. 6j). However, mCREB expression bidirectionally altered sensitivity to sunk costs. mCREB expression in the mPFC increased sensitivity while mCREB expression in the NAc decreased sensitivity to how much added value accumulates when experiencing irrecoverable losses while waiting. The opposing direction of these changes between mPFC and NAc mCREB treatment in wait zone sunk cost behavior are aligned with the opposing direction of changes in offer zone VTE behavior for negatively valued offers as well as sensitivity to the effect of type II violations on subsequent trials. This is in contrast to the shared direction of changes in VTE behavior for positively valued offers and sensitivity to the effect of type I violations on subsequent trials following mCREB treatment in either the mPFC or NAc. Groups did not differ in the way they valued other time spent on this task (Fig. S8). Taken together, these data provide a rich neuroeconomic framework to dissect differential region-specific roles of CREB in regulating complex decision-making computations across the mPFC and NAc that suggest a link between the value-based processing of offer zone and wait zone choices to sensitivity to distinct types of economic violations on future choices.

### Modeling the economic utility of sensitivity to distinct violations

In order to better understand the contribution of sensitivity to distinct economic violations on the overall ability of the individual to forage effectively for rewards, we generated a computer model of the Restaurant Row task that could accurately simulate mouse performance (Fig. 7a-b, Fig. S9). Because the Restaurant Row task involves multiple, complex interacting decision steps, it can be hard to predict how one decision variable affects the way animals forage throughout the rest of the session. By passing in each animal’s average decision speed, travel speed, reward consumption speed, and thresholds reflecting subjective flavor preferences, we could reliably simulate sessions of the task after presenting randomly generated sequences of offers. Thus, this Restaurant Row simulation offers the ability to systematically manipulate key decision variables and observe their downstream effects on performance outcome measures such as total number of end-of-session rewards earned. This simulation incorporated two regret terms called “type I bias” and “type II bias” that carried added value promoting the acceptance of negatively valued offers during subsequent decisions following each type of economic violation. Importantly, this regret term alters not the frequency with which mice make economic violations but rather to what degree violations influence the subsequent trial. These two regret terms, as well as violation rates, were systematically varied and revealed complex interactions influencing the relative number of total rewards earned (Fig. 7c, Fig. S10). We found that increasing sensitivity to type I bias resulted in an overall relative decrease in number of rewards earned on this task simulation (Fig. 7c). This was true across several variations of threshold violation rates (Fig. S10a), even if violation rates were different in each restaurant (Fig. S10b). Increasing sensitivity to type II bias affected earn potential to a much lesser degree (Fig. 7c, Fig. S10). We next examined at a deeper level if differences in regret terms among each restaurant could result in relative changes in flavor-specific reward earnings. We set sensitivity to each type of regret separated by either the least preferred (LP) or most preferred (MP) restaurants to match the profiles of CON, SUS, and RES mice (Fig. 7d, Fig. S10c-e). These simulations were compared against 1,000 shuffled control simulations that randomized the bias weight assignment from 0 to 1 across both types of regret and the different restaurants (Fig. S10c-d). Interestingly, sensitivity to type II bias in the LP restaurant, reminiscent of CON, RES, and GFP-only treated mice, resulted in no net change in reward intake in either LP or MP restaurants (Fig. 7d, Fig. S10e). Conversely, sensitivity to type I bias in the MP restaurant – the economic phenotype reminiscent of SUS and NAc-mCREB treated mice – resulted in the greatest change in reward intake (Fig. 7d, Fig. S10e). Furthermore, this economic phenotype resulted in a net positive gain in reward intake for LP flavors and a net negative loss in reward intake for MP flavors. These data suggest that distinct regret-related processes may have different downstream consequences on net foraging behavior. These data also suggest that the pattern of how mistake history of SUS mice influences future decisions may contribute to a redistribution of reward value shifted away from preferred rewards.

**Figure 7.**
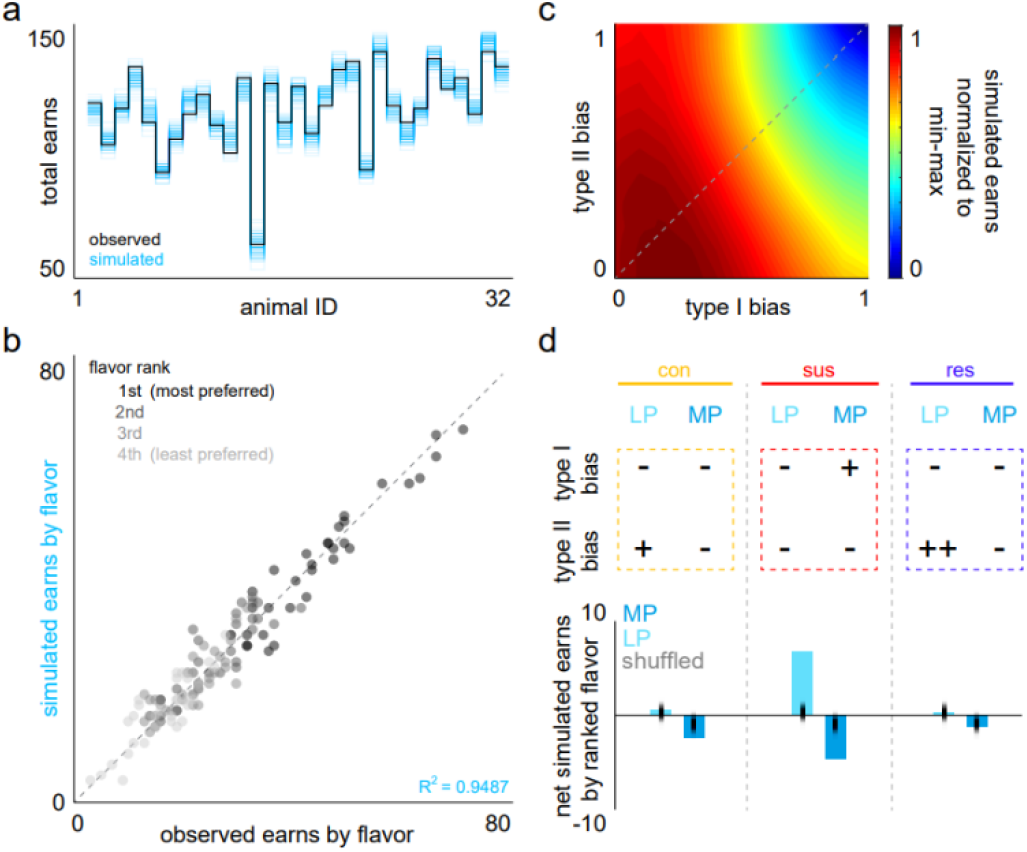
Economic modeling of regret-related phenotypes reveals different shifts in value-based reward intake. (a) Simulation of the Restaurant Row task quantifying total number of end of session earns, run 500 times (blue traces), in unique computer-generated sessions with randomly selected offers compared to observed earns (black trace) from an example session displayed across the 32 mice from the social defeat cohort. (b) Scatter plot of observed and simulated earns split by restaurant ranked by subjective flavor preferences with a high degree of concordance. See Fig. S9 for simulation data compared to the CREB cohort. (c) Simulations computing relative number of total pellets earned by varying the weight of type I violations on inflating the value of subsequent offers along the x-axis and the weight of type II violations on inflating the value of subsequent offers along the y-axis. Origin (0,0) indicates simulations ran where neither type of violation impacted subsequent decisions and individual trials were effectively treated as independent events. Coordinate (1,1) indicates simulations ran where both types of violations biased animals to always enter negatively valued offers on the subsequent trial. Note: when type I bias is set to 0, change in sensitivity to type II bias has a relatively little effect on net earnings, yielding overall more rewards relative to increases in sensitivity to type I bias. See Fig. S10 for simulations run with varying levels of threshold violation rates. Violation rates used in the simulations in this figure were set at 0.1, which approximate observed violation rates and yield smallest difference between simulated and observed data. (d) Difference between simulated earns and average observed earned in the least preferred (LP) and most preferred (MP) restaurants by setting bias weights independently in each restaurant to match the economic profile of CON, SUS, and RES groups compared against 1,000 simulations randomly shuffling bias settings. See Fig. S10 for the full matrix of bias weights organized by restaurant.

## Discussion

The way an individual experiences regret may be altered in stress-related disorders such as depression (*8*). However, this is often clinically described using plain language that, without attention to neuroeconomic principles, may miss fundamentally distinct computations derived from separable brain functions (*8, 10*). Here, we reveal two dissociable forms of regret-related behaviors in mice that are differentially associated with unique stress-response traits and CREB function in two brain regions (Fig. 8). Our findings suggest that the ability to appraise one’s own mistakes is made up of multiple processes that can be altered independently. These data have important implications for better understanding how different behavioral responses to poor decisions may be linked to adaptive versus maladaptive responses to stress.

This study revisited findings from Steiner and Redish 2014 and Sweis et al. 2018 to take a closer look at discrete action-selection processes involved in distinct economic violations compared to what an individual could have done differently (*5, 7*). Through complex analyses of behavioral sequences, we were able to measure differential weight animals must be placing on previously unselected actions during specific economic situations. We achieved this by looking at the influence of decision history on subsequent choices. To better describe the phenomena studied here, we turn to the decision psychology literature for more precise language surrounding the concept of regret. The nature of an unselected action and their outcomes can vary in several ways. Two dimensions along which to classify a counterfactual outcome include (i) direction and (ii) operation (*28, 29*). The direction of a counterfactual outcome may be either upward or downward. An upward counterfactual comprises one that would have been better than the actual outcome whereas a downward counterfactual comprises one that would have been worse than the actual outcome. Regret by definition stems from unselected actions that derive from upward counterfactuals and is what separates regret from simpler outcome evaluation or reward prediction error processes. The operation of a counterfactual outcome may be either additive or subtractive (*28, 29*). An additive counterfactual describes the unselected option that an individual recognizes is something one could have opted in to act on but actually did not. Conversely, a subtractive counterfactual describes the unselected option that an individual recognizes is something one could have forgone but actually acted on. In this study, economic violation type I would most likely evoke representations of an additive counterfactual. After rejecting an economically advantageous offer when subsequently encountering a worse offer, an individual may represent the missed opportunity of not accepting the previous offer. Here, the operative phrase is “not accept” that describes the erroneous nature of the actual outcome and thus emphasizes the counterfactual alternative for which one could have instead opted into by adding an action. Economic violation type II on the other hand would most likely evoke representations of a subtractive counterfactual. After accepting an economically disadvantageous offer and during corrective change-of-mind decisions, an individual may represent the alternative scenario in which the erroneous action could have been subtracted from reality thus negating the need to change one’s mind. While this is just one interpretation of the computations that may be at play during the behaviors measured here, it is nonetheless a useful framework for more easily describing the distinctions between the types of economic violations operationalized here and for making explicit predictions of the neural representations underlying different forms of counterfactual thinking.

Several examples of neural representations of counterfactual processing have been previously demonstrated across species. In humans, activation in medial subregions of the prefrontal cortex correlate with negative discrepancies between actual and counterfactual outcomes on gambling tasks and with self-report of the experience of regret only when information is provided about the outcome of unselected actions (*30*). Patients with damage to these brain regions reveal an inability to process and consider anticipated regret during decision making (*31*). In non-human primates, prefrontal neurons can encode hypothetical outcomes and reward signals that contribute to fictive learning with sustained changes in activity leading into subsequent decisions depending on whether or not information about the optimal choice is provided to the animal (*3, 32*). More recently, prefrontal and striatal neurons in non-human primates have been shown to encode the value of counterfactual outcomes of unselected actions when presented with the opportunity to select one of two reward offers presented serially regardless if the first or second of these two options were selected (*33, 34*). While such studies have begun to lay the foundation for how counterfactuals may be encoded in the brain and how they depend on access to information about the outcome of one’s actions, the neural underpinnings of both the direction and operation of regret-related processes and how this could map on to individual trait differences has not yet been explored prior to the present study.

In previously published work on the Restaurant Row task in rats, Steiner and Redish found that frontal and striatal ensembles could decode information about the missed opportunity on trial t-1 during the read-out trial (*5*). This was true only upon being informed of the subsequent high cost offer following economic violation type I but not during control sequences. These ensembles represented not the missed reward but rather the unselected choice (*5*), suggesting that counterfactual thinking evoked by type I violations may be more action- as opposed to reward-centric. It is surprising that in the present study, no control animals (non-defeated or GFP-only treated mice, in addition to RES mice) displayed sensitivity to this type of economic violation during subsequent decisions. This may be due to important task differences between that employed by Steiner and Redish and the present study. The task variant used by Steiner and Redish effectively had no offer zone – all offers immediately began counting down upon restaurant entry. Therefore, no explicit enter decision was made in that task variant as was measured in the present study. Thus, all skip decisions measured by Steiner and Redish also constituted quit decisions with no ability to clearly identify ballistic, erroneous enter decisions or distinguish economic violation type I from type II without an animal earning food on trial t-1, adding more confounds. Alternatively, and perhaps more interesting, is the possibility that the regret-related processes captured by Steiner and Redish indeed only track with those found here in SUS or mCREB-treated animals. Thus, close attention should be paid to animals who may have baseline elevated levels of stress either as a result of a species or strain difference or as a result of stressful laboratory experiences such as maintaining chronically implanted tetrodes. Nonetheless, we highlight here the importance of task design in being able to break down computations into discrete, measurable units.

Sensitivity to different types of regret may be linked to differences in the decision-making processes measured in offer zone and wait zone on trial t-1. VTE behavior measured in the offer zone is thought to reflect underlying deliberation between competing alternatives (*17-19, 35, 36*). For instance, during VTE, hippocampal place cell sequences sweep ahead of the animal along alternating paths between potential goals and correlate with reward representations in the mPFC and NAc (*4, 19, 37*). These data suggest that animals are evaluating the predicted outcomes of future options serially. Neural activity in both rodents and humans tested on translated versions of the Restaurant Row task indeed show signs of planning during offer zone decisions (*25-27, 38*). Here, mice treated with mCREB in the mPFC or NAc showed (i) shared changes in VTE behavior (decreased) when processing positively valued offers that covaried with sensitivity to regret type I (increased) as well as (ii) bidirectional changes in VTE behavior when processing negatively valued offers that covaried with bidirectional changes in sensitivity to regret type II. These data suggest that the decision process involved in the choice on trial t-1, reflected through VTE behavior, may be linked to the different value states the animal may be in upon arrival at the subsequent restaurant, differentially contributing to regret sensitivity. Others have reported that an attentional drift-diffusion model can explain how value-based decisions are made as reward value accumulates while individuals alternate attending to competing options with value signals that track in the prefrontal cortex (*39*). Furthermore, changes in VTE may reflect the level of indecisiveness as others have shown that value-based choices depend not only on past rewards and present confidence but also past confidence (*25, 40-43*). For instance, activity of both mPFC as well as midbrain dopamine neurons correlate with the predicted value of a chosen option that is the product of decision confidence and past value. However, only mPFC activity is causally linked to pre-outcome evaluations during on-going choices while dopamine activity is causally linked to evaluations only after a decision has been made and at a time when their prediction error signals are graded by confidence measures. Chemogenetic disruption of mPFC activity during Restaurant Row can decrease VTE, decouple mPFC and hippocampus interactions, and indirectly alter hippocampal sequences, causing animals to become more decisive (26, 38). Therefore, alterations in either of these constructs – value accumulation and decision confidence – may point to different neurophysiological changes during VTE on trial t-1 that could affect how mistakes bias future decisions, aspects of which may be uniquely altered in SUS versus RES mice.

Reward-related and stress-related phenotypes of SUS and RES mice have been well-characterized across a number of relatively simple behavioral screening tests (*15*). Exposure to stress increases CREB activity in the NAc (*14*). Elevated CREB function in the NAc reduces the rewarding effects of sucrose, consistent with an anhedonia-like response (*44*). Elevated CREB levels in the NAc also reduce responses to aversive stimuli, thought to be contributing to a generalized numbing of behavioral responsivity (*14*). Conversely, lower CREB function in the NAc has been linked to heightened behavioral responses to both rewarding and aversive stimuli (*11, 14, 45, 46*). CREB function in the mPFC is less understood, however, has recently been shown to promote RES phenotypes, as CREB knockout in the mPFC promotes SUS phenotypes (*13*). Although different from CREB knockout, in the present study CREB knockdown in the mPFC mimics the gain of sensitivity to regret type I, like SUS mice, but simultaneously mimics enhanced sensitivity to regret type II, like RES mice (*13*). Thus, our neuroeconomic findings partially overlap with a classic understanding of SUS versus RES mouse behavior. Because animals may use different circuits to achieve seemingly similar computations when tested on simple tasks, previous studies may have been unable to attribute which computations unique to a given brain region might be differentially perturbed in these mice (*47*). For instance, blunted dopamine signals in the NAc in contrast to enhanced dopamine transients in the mPFC describe known physiological signatures of SUS mice that could differentially encode the valence of error detection following type I and type II violations. Furthermore, the downstream consequences of CREB action can yield diverging cellular events that may govern separable domains of stress-related or reward-related behavior, even within the same brain region (*11*). Close attention to complex decision processes should be paid in translational research on depression in order to fit sometimes conflicting basic science findings and clinical symptomologies within a broader unified framework. For instance, although changes in reward prediction error signals seen in depressed patients can be divorced from mood symptoms (*48*), how individuals assign credit to internal errors during counterfactual thinking may be multifactorial. In a version of the Restaurant Row task translated for use in humans, risky decisions with poor outcomes, akin to regret type I, activated the default mode network and elicited altered behavioral responses in individuals with highly externalizing traits such as negative urgency, or the tendency to act rashly when distressed, that have been linked to depressive personalities (*27, 49, 50*). Such computations may elicit a negative bias when receiving late-arriving information about the optimality of a choice during regret type I sequences that generate a reactive response of being “let down,” and could reflect enhanced negative self-blame sometimes seen in depression (*9, 51-53*). This may be distinct from insensitivity to regret type II during self-initiated and volitional change-of-mind decisions which could stem from a failure to integrate self-monitoring and counterfactual representations with re-evaluating recent procedural mistakes (*21*). Thus, increased negative affect could drive enhanced sensitivity to regret type I independent of how anhedonia could blunt sensitivity to regret type II simultaneously within the same individual.

Although SUS and RES phenotypes are generally considered to be maladaptive and adaptive stress-response traits, respectively, it remains unclear whether or not sensitivity to regret type I or type II comprise maladaptive versus adaptive decision-making processes themselves. Unpacking this question is three-fold: (i) Do different regret-related processes impact foraging efficiency? (ii) Do different regret-related processes impact emotional burden? (iii) What is the validity of the rapid social interaction screening assay in separating SUS and RES mice into maladaptive versus adaptive traits? (i) First, to address the effect of decision bias conferred by each type of economic violation on food intake, we created a toy model of the Restaurant Row task that allowed us to systematically vary sensitivity to regret type I and type II while holding all other behavioral variables constant. We found that the regret-related economic phenotype unique to SUS mice produced the greatest change in food intake, but that this change interestingly resulted not in less overall food earned but rather a redistribution of yield shifted away from most preferred flavors and toward least preferred flavors. This analysis suggests that this phenotype indeed has a consequence on foraging efficacy that at face value appears to be detrimental to the individual and may add nuanced characteristic to more complex computations underlying outwardly appearing anhedonia-related behavior. (ii) Whether or not each type of regret differentially contributes to an affective component of mistake processing is more difficult to address here and may be unrelated to food intake altogether. Future animal studies should compare individual differences in sensitivity to each type of regret with propensity to demonstrate other affect-related processes, including startle, anxiety-like behavior, and fear conditioning, for example. Furthermore, because this task has been translated for use in humans (*22, 27, 49, 50, 54*), accessing emotional states immediately after each decision step would be useful in better approximating these constructs across species and in patient populations. (iii) A larger question for the field is whether or not the rapid social interaction screen is a valid marker for identifying stress-suppressibility versus stress-resilience (*55*). This assay has served as a useful tool for predicting numerous other depressive-related phenotypes characterized on a wide battery of tasks that have served as the basis for the development of much of the pharmacological treatments for depression (*12, 15*). RES mice defined by this assay too reveal several gene expression changes including in CREB beyond that of control mice that are casually linked to antidepressant phenotypes and map on molecular fingerprints in human post-mortem tissue that is lost in patients diagnosed with depression (*13*). Nonetheless, some have challenged this notion of the adaptiveness of SUS and RES mice in several different ways. For instance, social avoidance demonstrated by SUS mice can be considered to be adaptive and could promote survival in the face of perceived threat (*56*). Recently, others have attempted to characterize the cost-benefit trade-off of different stress-related traits among SUS and RES mice that may have more nuanced implications for passive versus active coping styles, social discrimination skills, as well as physiological stress responses measured peripherally (*57*). It remains an ongoing endeavor to explore how other classification systems may be able to characterize individual differences in stress-related disorders. Taken together, we nonetheless demonstrate here for the first time that distinct regret-related processes are indeed separable and map on to individual differences in stress-response traits. Of note, we report here for the first time a behavioral difference between RES mice and non-defeated CON animals apart from that of SUS mice. RES mice revealed enhanced sensitivity to regret type II as well as, related, enhanced sensitivity to sunk costs during change-of-mind decisions. It is interesting to point out that sensitivity to sunk costs in particular has historically been thought to be part of an economically suboptimal decision-making bias, which brings into question why such a phenomenon has been conserved across evolution (*22, 58-60*). A rational agent, in theory, should ignore sunk costs when making economic decisions. Economic stress, budget constraints, and limited reward availability indeed can drive individuals to make suboptimal choices, for example, in the form of overly perseverative behaviors that can reallocate finite resources sometimes with diminishing returns (*24, 61*). Sensitivity to sunk costs is known to be heightened under such circumstances (*22, 23*). However, it has been postulated that sensitivity to sunk costs may have hidden utility (*22*). For instance, valuations calculated from predictions of future outcomes can be difficult and so basing decisions instead on past information may sometimes be a better predictor of future returns, which can serve as a useful heuristic when foraging (*24, 25, 59, 60, 62-65*). Alternatively, a sunk cost bias may emerge as a byproduct of enhanced sensitivity to change-of-mind-induced regret, which is typified by a gain in regret type II both in RES mice as well as mCREB-treated animals in the mPFC, and conversely reduced by mCREB expression in the NAc. Chemogenetic disruption of mPFC activity can drive an increase in sensitivity to sunk costs in Restaurant Row concurrent with changes in neural ensembles that represent more local and less forward-oriented information (*25, 26, 38*). Optogenetically depotentiating mPFC outputs to the NAc can also decrease the frequency of change-of-mind decisions (*66*). Collectively, these data suggest that RES animals may have evolved circuit-specific processes that rely on sunk costs and change-of-mind-induced regret to sharpen decisions and increase attention paid toward realized losses that may be more costly (*67*). It may also be possible that animals better suited to cope with stress-induced pathology, who also happen to engage decision valuations in this way even if unrelated to a stress response itself, may have indirectly contributed to why decision phenomena such as type II regret and sensitivity to sunk costs have been conserved across evolution (*68, 69*).

While this body of work represents a unique combination of elements from several different fields including stress models, molecular manipulations of transcription factors, and complex neuroeconomics, there are several limitations to this study. First, there are many different stress paradigms known to elicit heterogenous responses in animals. Here, mice were only tested in the chronic social defeat stress model of depression. Other stress protocols beyond social defeat, including chronic variable stress, early life stress, and chemical stress – protocols more easily suitable to probe potential sex differences – should be explored in future studies. Second, a common issue in stress-related research is understanding if behavioral effects stem from a set of pre-existing traits, are accentuated by stress, or are induced de novo following a stress exposure. That is, one concern is whether or not regret type I and regret type II may be characteristics present in SUS and RES mice prior to social defeat and prior to the designation of SUS and RES phenotypes as determined by the rapid social interaction screening test. It should be noted that the primary finding of this study disentangles each type of regret from one another, demonstrating that mistake appraisal does not simply generalize to a common computational basis for the monitoring of the error of one’s own agency but rather that these processes are indeed dissociable. A secondary finding is that these distinct processes independently covary in animals with individual differences in unique stress-response traits. How much one’s sensitivity to each type of regret may serve as a predictive tool for susceptibility versus resilience to stress is an interesting, although separate question. Nonetheless, one piece of evidence suggesting regret sensitivity is not a pre-existing characteristic of stress-naïve animals is the fact that the presence of regret type I does not exist at baseline in non-defeated CON mice or GFP-only treated mice. Baseline sensitivity to regret type II on the other hand in theory could be used to in a future experiment to predict behaviors on a rapid social interaction screening test following defeat. In addition, future studies should investigate the degree to which these regret-related processes may be reversible in defeated animals either through CREB manipulations similar to those presented here or following other treatment with classic antidepressant pharmacology, for instance. These experiments, together with in vivo physiology and an investigation of downstream molecular consequences of CREB function, would help link gene expression changes to the electrophysiological signatures of counterfactual thinking and help broaden the scope and complexity of depression research.

In summary, we operationalized value across several dimensions and revealed how choice history impacts future decisions not only based on the framing of one’s past mistakes but also based on what the individual could have done differently. We provide a novel lens through which to stratify more complex decision-making computations and identify two fundamentally distinct forms of regret-related processes that may evoke different additive or subtractive counterfactuals linked to susceptibility versus resilience to stress. Importantly, we provide insight toward identifying therapeutic targets for further investigation as to which regret-related processes may need to be potentially restored (type II) versus ameliorated (type I) in the treatment of stress-related disorders like depression. Our neuroeconomic approach to computational psychiatry has been validated in a set of tasks translated for use across species and affords a rich pipeline to directly apply discoveries from animal behavior to human psychology in ways that could provide new structure to how patients are interviewed clinically, asking specific questions about the nature of counterfactual thinking (*70*). This work demonstrates how circuit-computation-specific processes can be extracted based on a careful description of one’s decision-making processes and, in the case of the complexities of regret, about how not all mistakes are created equally and about the different roads not traveled.

## Acknowledgments

We thank members of the Nestler and Russo labs for helpful discussion and technical assistance. We thank Peter Rudebeck, Caleb Browne, Xaiosi Gu, Denise Cai, Antonia New, and Asher Simon for constructive conversation. We also thank Alexxai Kravitz for assistance in developing the open-source pellet dispensers used in this experiment (www.hackaday.io/project/171116-fed0). Open-source illustrations obtained from SciDraw (www.scidraw.io), credit Luigi Petrucco, Federico Claudi, Tiago Branco, and Gil Costa.

## Author contributions

Conceptualization: BMS; Methodology: RDC, SJR, EJN, BMS; Investigation: RDC, FMR, LL, AMT, FC, LMH, FY, SOE, JFS, SA, BMS; Data curation: RDC, BMS; Formal analysis: RDC, LMH, BMS; Funding acquisition: SJR, EJN; Supervision: RDC, SJR, EJN, BMS; Writing – original draft: RDC, BMS; Writing – review & editing: all authors

## Data and materials availability

All data, code, and materials used in the analysis are available in the materials and methods section or upon request. Supplementary materials available online.

## Materials and methods

### Animals and husbandry

10-week-old wild-type male C57BL/6J mice were purchased from Jackson Laboratory for the experiments in this study. Additionally, 16 to 24-week-old male CD-1 (ICR) mice (sexually experienced retired breeders purchased from Charles River Laboratories) were used as aggressors for the chronic social defeat stress (CSDS) protocol. All C57BL/6J mice were initially randomly group-housed (3-5 mice per cage) and allowed a 1-week period to acclimate to the housing facilities before the start of experiments. CD-1 mice were singly housed. During the CSDS protocol, a single C57BL/6J mouse was co-housed with a single CD-1 mouse as part of the CSDS protocol described in detail below. Prior to and during the CSDS protocol, mice had access to regular chow ad libitum. Following the CSDS protocol, C57BL/6J mice were individually housed and switched to a full-nutrition flavored pellet diet (BioServe products; 20 mg dustless precision pellets; a ∼3 g mixture of chocolate, banana, grape, and plain flavored pellets as a daily ration) and food restricted to approach 80-85% of their free-feeding body weight over the next 3 days before starting training on the neuroeconomic operant decision-making paradigm termed “Restaurant Row” described in detail below where mice work for these very same 20 mg rewards as their sole source of food. Mice were weighed daily before and during the CSDS protocol and twice daily (before and after testing) on the Restaurant Row task. Mice were placed daily in Restaurant Row 7 days a week from to the beginning to the end of the experiment in order to maintain the closed-economy contingency wherein task performance each day provided full nutrition. All mice were maintained on a 12-hr light/dark cycle with ad libitum access to water. Experiments were conducted during the light phase. In the rare instance (<5%) that mice could not independently support their own body weight by foraging for rewards on the task, small rations of post-task supplementary feeding were offered to the animals that they readily consumed before fasting again for 23 hr until the next Restaurant Row testing session to remain in concordance with animal safety regulations. Experiments were approved by the Mount Sinai Institutional Animal Care and Use Committee (IACUC; protocol number LA12-00051) and adhered to the National Institutes of Health (NIH) guidelines.

### CSDS

Mice underwent CSDS, a well-established animal model of psychosocial stress that is capable of inducing a depressive- and anxiety-like phenotype (28). CD-1 mice were screened for aggressive behaviors prior to use. During CSDS, a single C57BL/6J mouse was co-housed with a single CD-1 mouse and allowed to physically interact and experience aggression behavior for 5-10 min of attacking before being separated for the remainder of the day. Both mice remained co-housed in the same cage but separated by a mesh divider so the mice no longer had direct physical contact but continued to have visual, olfactory, and auditory contact. This procedure was repeated for 10 consecutive days with 10 different CD-1 mice. As a control to this stressor, non-stressed (non-defeated) C57BL/6J mice were handled equally without exposure to CD-1 mice but were exposed to other, domiciled C57BL/6J mice of the same size and age. By the end of this protocol, mice are typically assayed on a social interaction test that captures social avoidance, which has been shown to be highly predictive of several other depressive-like behavioral and neurobiological abnormalities (28). This assay is a short behavioral screen where a single C57BL/6J mouse is placed in a large open field arena with a novel CD-1 mouse enclosed in a small chamber. EthoVision software was used to track the location of the C57BL/6J mouse during this social interaction assay. Time spent near (interaction zone) versus away from the CD-1 mouse was used to quantify a social interaction score calculated from time in the interaction zone with the CD-1 mouse present in the chamber (2.5-min trial) relative to a preceding 2.5-min baseline trial without the target CD-1 mouse present.

### Neuroeconomic Decision-Making Paradigm: Restaurant Row Task

Mice were trained to forage in a square maze for food rewards of varying cost (delays ranging from 1 to 30 s) and subjective value (unique flavors tied to four separate and uniquely spatially cued locations, or “restaurants,” located in each corner of the maze) while on a daily limited time budget (60 min). Each restaurant consisted of two separate decision zones, (1) an offer zone (T-shaped intersection) and (2) a wait zone (small chamber with a reward receptacle). Upon entry into a restaurant’s initial offer zone, a tone sounded whose pitch indicated the delay mice would have to wait in order to earn food if they chose to enter the wait zone (higher tone pitch equates to a longer delay randomly selected upon offer zone entry; pitch identities were shared across restaurants). If a mouse chose to enter the wait zone, a countdown began during which tones descend stepwise every second either until the reward was earned or the mouse decided to quit and leave the wait zone. There is no penalty to quitting other than the offer was rescinded and the mouse must advance to the next restaurant. Thus, a trial was terminated if a mouse made: (1) a skip decision in the offer zone and advanced down the hallway, (2) a quit decision in the wait zone, or (3) earned a reward, after which the mouse must progress to the next restaurant in a serial order. Importantly, rewards earned on this task served as the only source of food (full nutrition flavored BioServe 20 mg pellets), making this task closed-economic in nature with time as a limited commodity. This means that decisions made on this task were interdependent both across trials as well as across days. This also means that any time spent engaged in a given behavior was at the expense of spending from the time budget engaged in other behaviors or exploring alternative options, a concept known as opportunity cost in the foraging literature. Thus, how value is calculated can be operationalized in many forms depending on available actions to choose from, choice history, and current economic situation. Different animals preferred the unique flavors differently and such idiosyncratic differences in flavor preferences were harnessed to operationalize value subjectively as a function of offer cost relative to indifference points in decision thresholds within a given restaurant as well across the ordinal rankings of each restaurant’s flavor. Flavor preferences developed early in training and were stable across days—roughly equal fractions of mice displayed preferences for each of the flavors. Each restaurant remained spatially fixed in the maze with patterns on the wall to signify the restaurant identity (chocolate: vertical stripes; banana: checkers; grape: triangles; plain, horizontal stripes.

Rewards were delivered using a 3D printed automated pellet dispenser that was triggered by a computer running the behavioral task programmed in the ANY-Maze software made by the Stoelting Company. Behavioral events were triggered by spatial movements through the maze tracked by ANY-Maze. The receptacle of the pellet dispenser also featured a custom-built trap door that would discard an uneaten pellet triggered upon exit from the wait zone if mice did not immediately consume food off of the pedestal. This prevented mice from hoarding rewards and quickly trained animals to adhere to the structure of the task to make meaningful and intentional foraging decisions. Small wall mounted speakers (MakerHawk 3 Watt 8 Ohm Single Cavity Mini Speakers driven by a DROK 5W+5W Mini Amplifier Board PAM8406 DC 5V Dual Channel Class D) were fixed to the wall of each restaurant that played a 500 ms tone upon entry into the offer zone and repeated every s until either an enter or skip decision was made. The pitch of the tone varied depending on the randomly selected offer of that trial (1 s = 4,000 Hz and each second above that was an additional 387 Hz; e.g., 5 s = 5,548 Hz; 15 s = 9,418 Hz; 30 s = 15,223 Hz). Upon entry into the wait zone, the tones descended in a countdown fashion stepping down 387 Hz each second until mice either quit the wait zone or waited out the full countdown. ELP USB camera with a Xenocam 1/2.7” 3.6 mm lens was used for video tracking. Restaurant Row testing took place in dim lighting conditions.

### Stereotaxic surgery and viral gene transfer

In a separate cohort, an additional 40 C57BL/6J mice (10 weeks old from Jackson Laboratories) were anesthetized by intraperitoneal injection with a mixture of Ketamine HCl (100 mg/kg) and Xylazine (10 mg/kg) and positioned on a stereotaxic instrument (David Kopf Instruments). In the nucleus accumbens (NAc, from bregma with an angle of 10 degrees: AP +1.6 mm; ML ±1.5 mm; DV −4.4 mm) or medial prefrontal cortex (mPFC, from bregma with an angle of 15 degrees: AP +1.8 mm; ML ±0.75 mm; DV −2.7 mm), 0.7-1 μL of virus (AAV2-CMV-mCreb, 1×1012 Addgene Plasmid #68551; or AAV2-CMV-eGFP, 1×1012 UNC GTC Vector Core) was bilaterally infused using 33-Gauge Hamilton needles over 5 min, and the needle was left in place for 5-10 min after the injection. Mice were allotted two weeks to recover before beginning food restriction in preparation for testing on Restaurant Row to match the timeline of the CSDS cohort. At the end of the behavioral testing animal were euthanized and viral transfection was visually inspected using a fluorescence microscope. Brain tissue used for histological quantification of virus transfection levels and cell-type specificity was collected in a separate set of test mice three weeks post-surgery. At time of collection, animals were deeply anesthetized with peritoneal injections of 500 mg/kg of Fatal Plus (Vortech, Cat #9373), and intracardially perfused with 15 mL 4% PFA (Electron Microscopy Science, Cat #15713-S). Brains were postfixed for 24-72 hr, and subsequently sliced on a Leica VT1000 S vibratome at 40-50 μM sections. Sections were blocked for 1 hr in blocking buffer (10% donkey serum (Jackson Immunoresearch, Cat #017-000-121), 0.3% Triton-X (Sigma, Cat #9284) in PBS), followed by overnight incubation with primary antibody (1:1000 Ch-NeuN, MilliporeSigma, Cat #ABN91) in diluted blocking buffer (1:3 dilution in PBS). Sections were washed three times with diluted blocking buffer (15 min each) before incubation with secondary antibodies (Ch-647: Jackson Immunoresearch, Cat# 703-605-155) for 1 hr. Two additional 15-min washes in diluted blocking buffer and one 15-min wash in PBS. Finally, sections were incubated with 1:10,000 DAPI (Thermofisher, Cat #62248) for 5 min. Sections were mounted with Prolong Diamond Antifade Mountant (Thermofisher, Cat #P36970). Images were acquired on a Zeiss LSM 780 confocal microscope using Zen software with 40x oil immersion lens at 1.1 digital zoom. Three images per region and animal were acquired. Quantification of transfection was performed using Cell Profiler. Cells were first identified to be DAPI+, neurons were subsequently identified if both DAPI and NeuN positive. Similarly, virally transfected cells were identified if both DAPI and GFP positive. Virally transfected neurons were identified if DAPI, NeuN, and GFP positive.

