## Supplementary Data for "Region-specific CREB function regulates distinct forms of regret associated with resilience versus susceptibility to chronic stress"

**This PDF file includes:**

Figs. S1 to S10

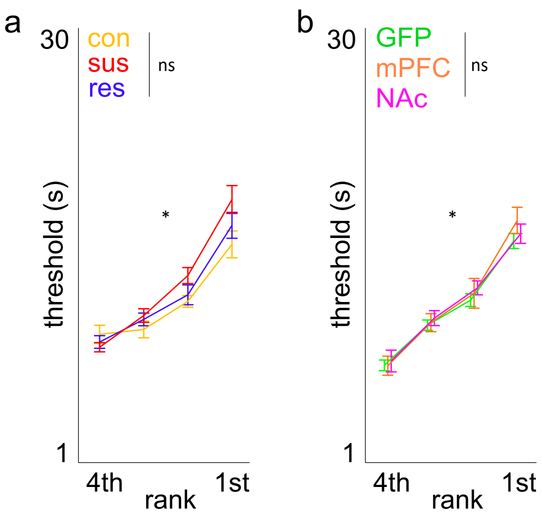


**Supplementary Figure 1. Economic thresholds of willingness to wait.** Thresholds were calculated by fitting a heaviside-sigmoid step function to choice outcome in each zone as a function of cued offer cost and identifying the inflection or indifference point of each curve. Thresholds were calculated daily and thus provide a way to normalize offer value across days as well as between restaurants across animals. Thresholds plotted split by restaurant ranked by subjective flavor preferences defined by end of session earnings for (a) social defeat cohort (significant main effect of rank: $F_{2,3}$=42.504, *$p$<0.0001; but no interaction between groups: $F_{2,3}$=0.775, $^{ns}p$=0.59) and (b) CREB cohort (significant main effect of rank $F_{2,3}$=52.175, *$p$<0.0001; but no interaction between groups: $F_{2,3}$=1.526, $^{ns}p$=0.17). Error bars ± 1 SEM. Not significant (ns).

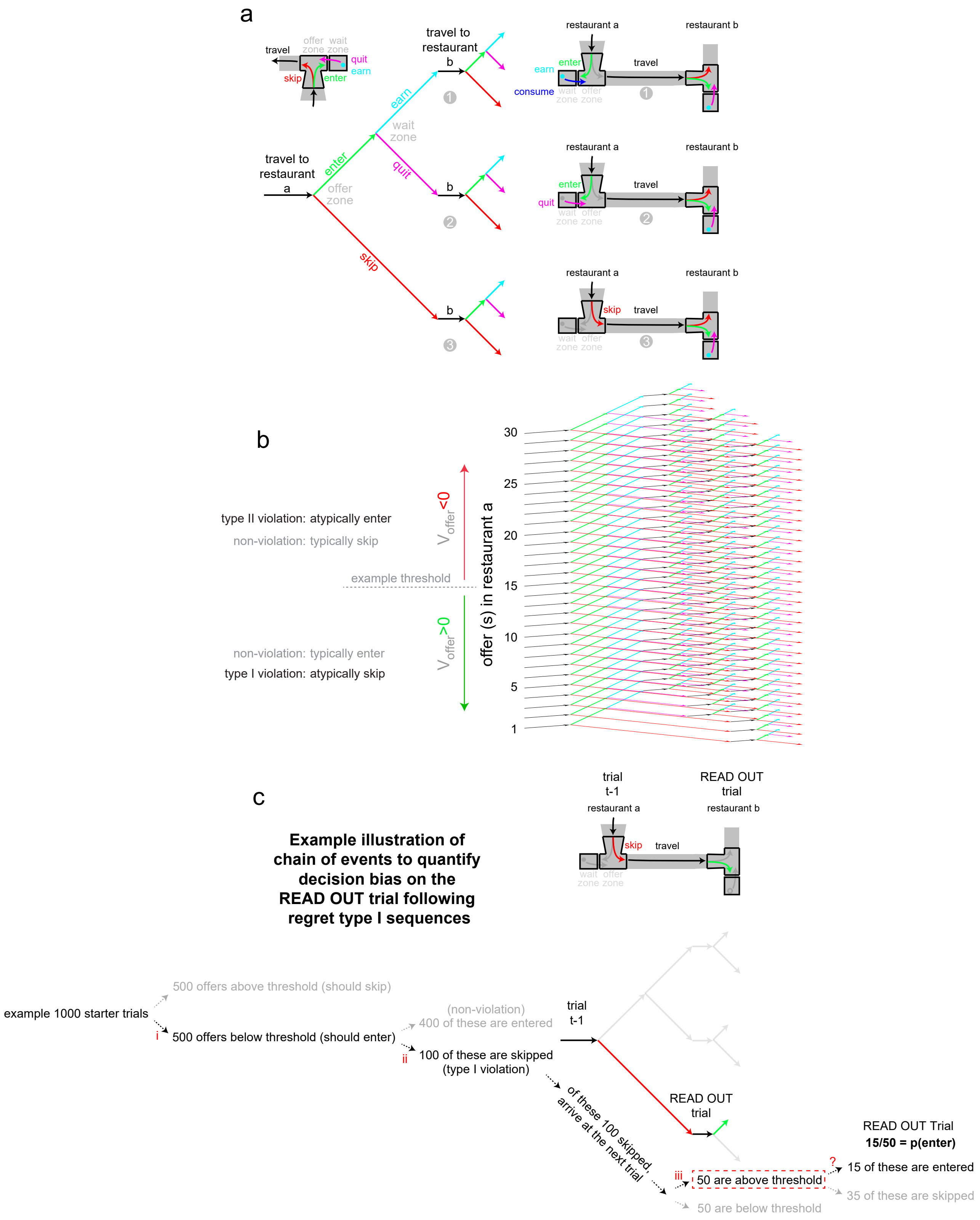


**Supplementary Figure 2. Visual explanation of regret-related analysis.** The three ways a trial on the Restaurant Row task terminates comprise: (1) earning a reward, (2) quitting in the wait zone, or (3) skipping in the offer zone. Examples of these trial-end outcomes are depicted in (a) via a decision tree (left) and maze schematics (right). These visuals also illustrate how trial sequences are constructed showing trial-end outcomes in "restaurant a" followed by a subsequent trial encounter upon arriving at "restaurant b." (b) Decision trees increase in complexity when factoring in what the initial offer of the starting trial in "restaurant a" is. For example, 30 decision trees are depicted here stacked atop one another. An example threshold of 15 s is indicated by the horizontal dashed gray line. Therefore, offer value [$V_{offer}$ = threshold – offer] is positive for all offers below this line and negative for all offers above this line. Atypical decisions, or economic violations, comprise trials in which mice skip offers below threshold (type I violation) or enter offers above threshold (type II violation). The complexity of these decision trees also increase based on several other conditional statements, for instance, including value contingencies of the offers in both "restaurant a" AND "restaurant b" as well as aligning trees to each restaurant ranked by subjective flavor preferences. (c) An example illustration of the sequence used to calculate the primary decision bias on the read out trial in "restaurant b" is depicted. This visualization is intended to show how the probability of accepting an offer on the read out trial is determined from a subset of "starter trials" in order to isolate the effect of choice history (for instance, violation versus non-violation choice history on trial t-1) on subsequent valuations. In this example, a subset of encounters on the read out trial (50, emphasized by dashed red box) satisfy the contingencies that go into the regret-related analysis for type I violations: (i) the offer on trial t-1 was positively valued, (ii) the choice on trial t-1 was a skip decision, and (iii) the offer on the read out trial was negatively valued. These contingency steps are labeled by red-colored "i, ii, and iii" at each arrow. From these remaining trials in this example (50, denominator), the number of encounters on the read out trial that result in an enter decision in the offer zone (15, numerator) are used to calculate the decision bias metric (emphasized by the red-colored "?"). This analysis is used for all other scenarios varying respective contingencies (i.e., non-violation events on trial t-1, for type II violations and its respective non-violation sequence, as well as additional control read out scenarios as described in Fig. S5).

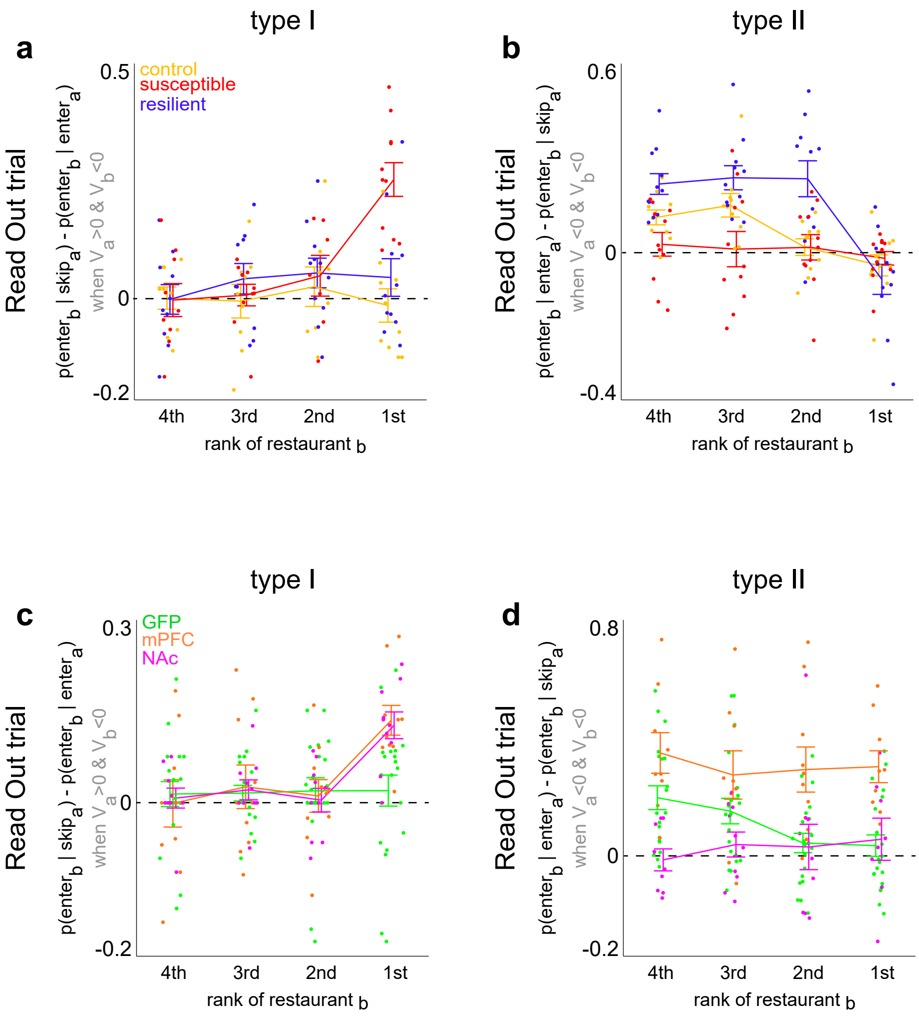


**Supplementary Figure 3. Difference score of decision bias on read out trials between violation and non-violation sequences.** Quantification of the magnitude of decision bias on read out trials. This effect stems from the difference between violation versus non-violation choices made on trial t-1, holding all other parameters constant and aligned to the ranking of the restaurant on the read out trial. Data plotted here comprise a difference score between the black (violation) and gray (non-violation) lines in Fig. 3b,d for the social defeat cohort (a-b) and Fig. 6e,g for the CREB cohort (c-d). This difference score describes a metric that captures decision bias, or change in the probability of entering negatively valued offers on the read out trial due to a violation choice history relative to a non-violation choice history. This metric operationalizes sensitivity to regret-related processes that can influence subsequent valuations. Type I regret-related bias plotted in (a, social defeat) and (c, CREB) and type II regret-related bias plotted in (b, social defeat) and (d, CREB). Horizontal dashed line reflects net change of zero and thus an insensitivity to a given type of violation choice history on subsequent valuations. Offer value is defined by [$V_{offer}$ = threshold – offer]. Y-axis nomenclature in panel (a) describes the difference score and is defined as follows: when the value of offer "a" is positive and when the value of offer "b" is negative, the probability of entering "restaurant b" given a skip in "restaurant a" (type I violation) minus the probability of entering "restaurant b" given an enter in "restaurant a" (non-violation). Other y-axes in this supplementary figure follow similar difference scores for type I and type II sequences accordingly with respective non-violation control sequences. ANOVAs: (a) $\text{interaction}_{\text{rankXgroup}}$: $F_{2,3}$=4.572, *$p$<0.001. (b) $\text{interaction}_{\text{rankXgroup}}$: $F_{2,3}$=4.140, *$p$<0.001. (c) $\text{interaction}_{\text{rankXgroup}}$: $F_{2,3}$=2.468, *$p$<0.05. (d) $\text{interaction}_{\text{rankXgroup}}$: $F_{2,3}$=2.432, *$p$<0.05. Dots represent individual animals. Error bars ± 1 SEM. Not significant (ns).

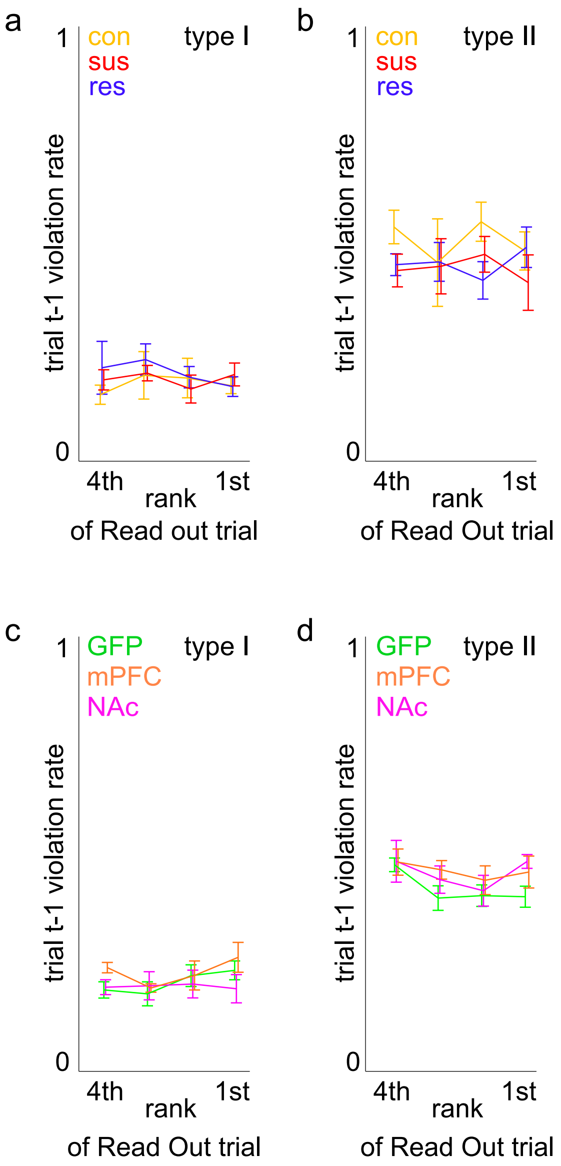


**Supplementary Figure 4. Threshold violation rates on trial t-1 split by restaurant aligned to the read out trial and ranked by subjective flavor preferences.** (a) In order to quantify the frequency of type I violation decisions made on trial t-1 aligned to the ranking of the restaurant on the read out trial, the probability of skipping a positively valued offer was calculated by identifying all positively valued offers encountered at the restaurant before the rank indicated on the x-axis (denominator) and identifying the subset of those trials that were skipped (numerator). There was no significant main effect of rank ($F_{2,3}$=0.689, $^{ns}p$=0.56) and no interaction by group ($F_{2,3}$=0.292, $^{ns}p$=0.94). (b) A similar analysis was performed for type II violations, identifying all negatively valued offers encountered at the restaurant before the rank indicated on the x-axis (denominator) and identifying the subset of those trials that were entered (numerator). (rank: $F_{2,3}$=2.354, $^{ns}p$=0.09; interaction by group: $F_{2,3}$=1.007, $^{ns}p$=0.43). Data shown for the (a-b) social defeat cohort and (c-d) CREB cohort (c, type I, [rank: $F_{2,3}$=0.640, $^{ns}p$=0.59; interaction by group: $F_{2,3}$=0.542, $^{ns}p$=0.77]) and (d, type II, [rank: $F_{2,3}$=1.730, $^{ns}p$=0.16; interaction by group: $F_{2,3}$=0.661, $^{ns}p$=0.68]). Error bars ± 1 SEM. Not significant (ns).

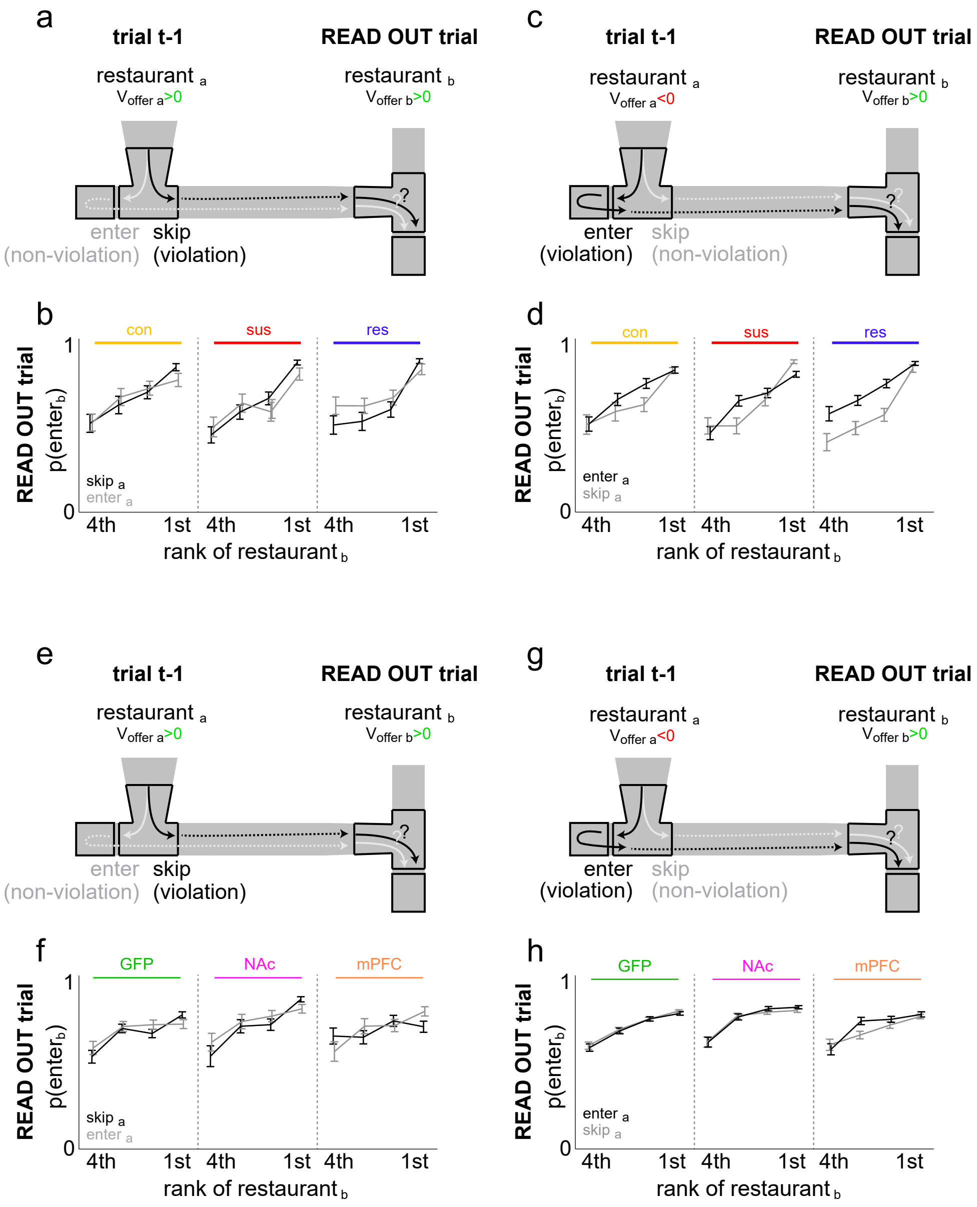


**Supplementary Figure 5. Economic violation sequences involving a control read out trial with positively valued offers presented instead.** Data for additional control sequences in which the read out trial presents a positively valued offer is depicted. All analysis in this figure follow similar patterns as described in Fig. 3. The effects of type I violations and respective non-violation sequences during trial t-1 on subsequent read out trial decisions shown in (a-b, social defeat cohort: two-way ANOVA group x choice history: $F_{2,1}$=2.350, $^{ns}p$=0.10) and (e-f, CREB cohort, $F_{2,1}$=0.371, $^{ns}p$=0.69). The effects of type II violations and respective non-violation sequences during trial t-1 on subsequent read out trial decisions shown in (c-d, social defeat cohort, $F_{2,1}$=3.827, $*p<0.05$) and (g-h, CREB cohort, $F_{2,1}$=2.136, $^{ns}p$=0.12). Error bars ± 1 SEM.

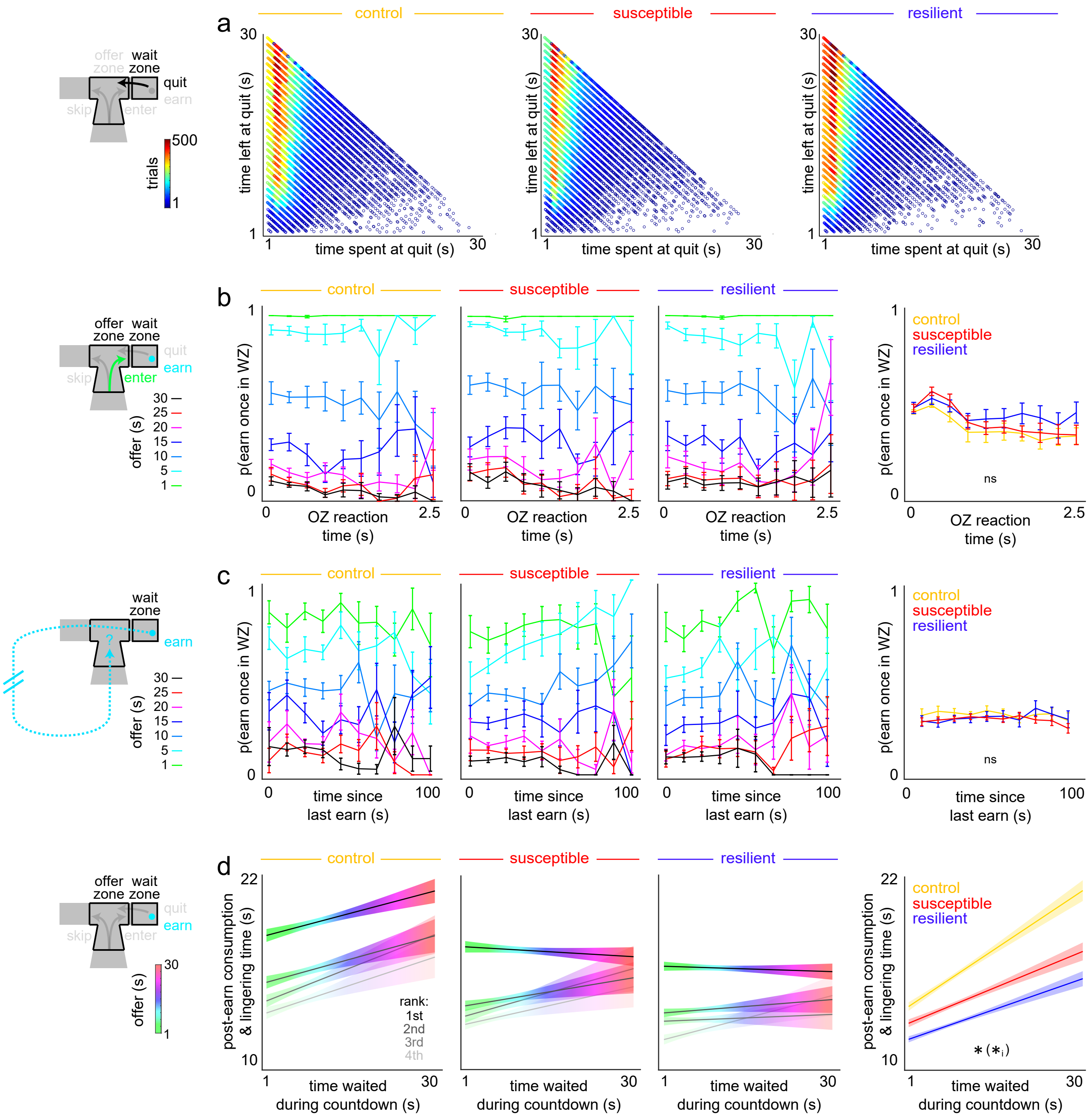


**Supplementary Figure 6. Quantification of how mice from the social defeat cohort uniquely value different forms of time spent on the Restaurant Row task.** (a) Quit trials pooled across all animals parsed into [time spent, time left] pairs measured at the moment of quitting relative to countdown onset or time to reward delivery, respectively. (b) The amount of time spent before making an enter decision in the offer zone (OZ) has no impact on the probability of earning a reward once in the wait zone (WZ), separated by offer (left) or collapsed across all offers (right, two-way ANOVA between group and OZ reaction time: $F_{2,9}$=1.657, $^{ns}p$=0.192). (c) The amount of time elapsed since the last pellet was earned has no impact on the probability of earning a reward once in the WZ, separated by offer presented upon arrival into the trial from which p(earn) is calculated (left) or collapsed across all offers (right, two-way ANOVA between group and time elapsed since last earn: $F_{2,9}$=1.601, $^{ns}p$=0.203). (d) The amount of time spent consuming and lingering at the reward site after earning a reward increases a function of time already waited during the countdown (i.e., offer length) split by ranked flavor preferences (left) or collapsed across restaurants (right). Significant main effect of time waited on post-consumption lingering time ($F_{29,31}$=75.090, $^{*}p$<0.0001) and a significant interaction with group ($F_{2,29}$=22.505, $^{*}p$<0.0001). The data in (d) suggest mice are differentially sensitive to time spent waiting in the wait zone, related to sensitivity to sunk costs during change-of-mind decisions. Error bars ± 1 SEM. Not significant (ns).

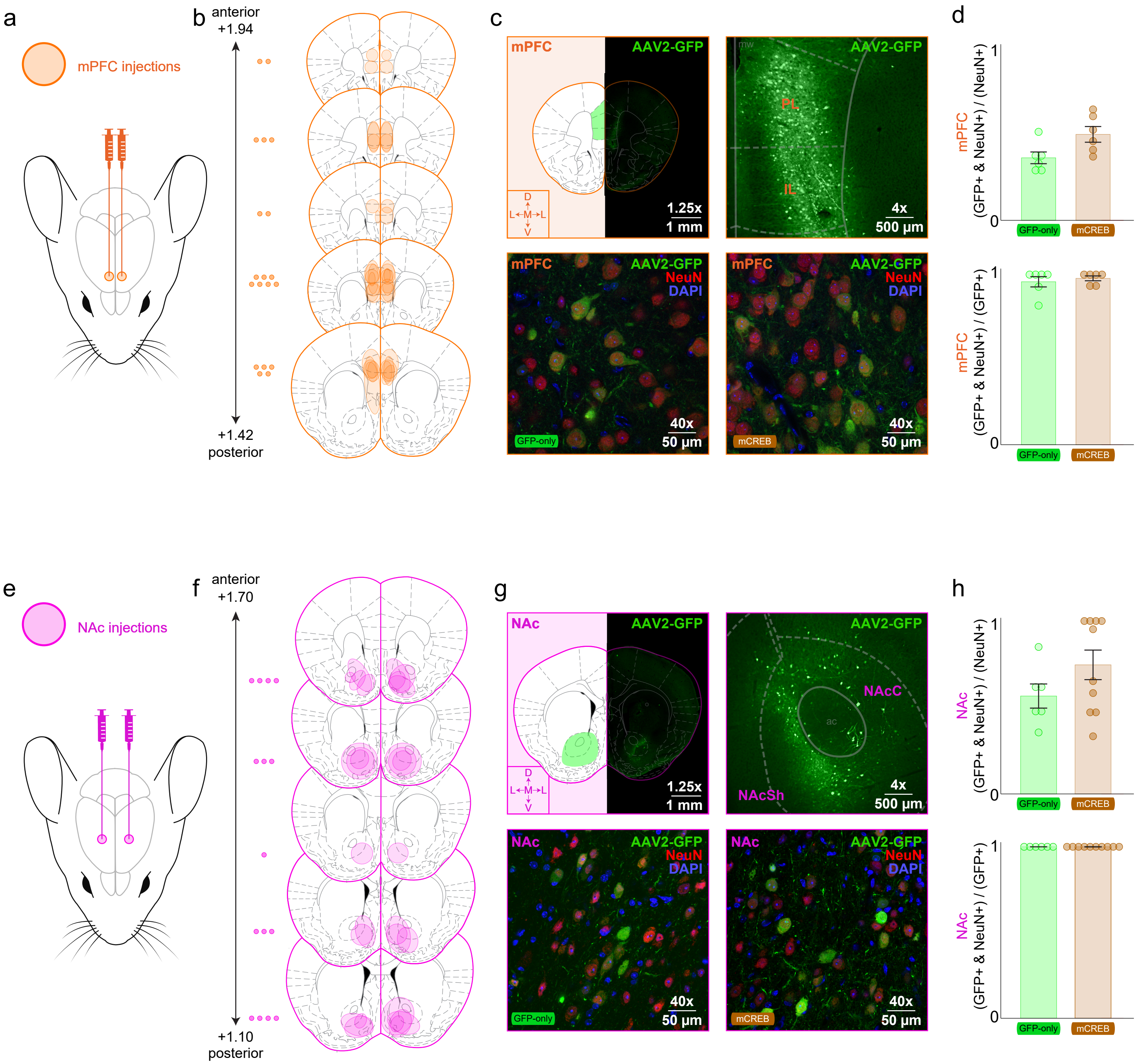


**Supplementary Figure 7. Virus targeting.** (a) Schematic of bilateral injection sites for medial prefrontal cortex (mPFC). (b) Anterior to posterior axis of coronal sections showing distribution of transfection targeting, coordinates in mm units relative to bregma. Dots represent individual animals. Overlapping shaded regions reflect spread of transfection. (c) Representative images of green fluorescent protein (GFP) reporter expressed in the adeno-associated virus 2 (AAV2) GFP-only or mCREB-containing viral construct co-labeled with nuclear DAPI and neuronal NeuN staining at 1.25x, 4x, and 40x. Inset depicts dorsal (D) / ventral (V) and medial (M) / lateral (L) axes. (d) Quantification of proportion of total neurons transfected (top) and proportion of transfected cells that are neurons (bottom). Note: even by using an AAV2 driven by a CMV promoter, we were able to achieve transfection essentially only in neuronal cell types. (e-h) Same as (a-d) except for targeting in the nucleus accumbens (NAc). Labels in 4x images: Prelimbic cortex (PL). Infralimbic cortex (IL). Medial wall (mw) of the prefrontal cortex. NAc Core (NAcC). NAc Shell (NAcSh). Anterior commissure (ac). Error bars ± 1 SEM.

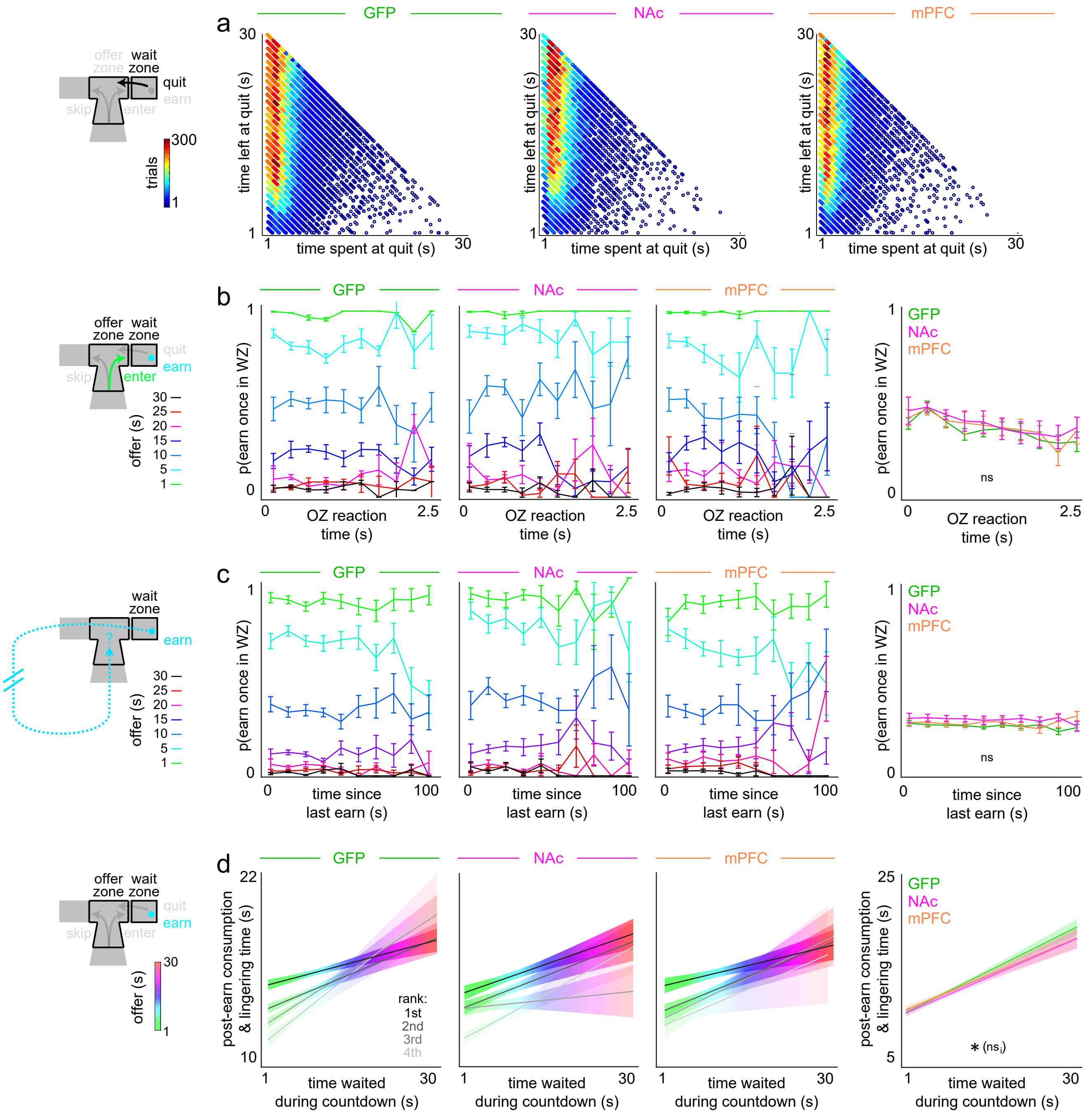


**Supplementary Figure 8. Quantification of how mice from the CREB cohort uniquely value different forms of time spent on the Restaurant Row task.** (a) Quit trials pooled across all animals parsed into [time spent, time left] pairs measured at the moment of quitting relative to countdown onset or time to reward delivery, respectively. (b) The amount of time spent before making an enter decision in the offer zone (OZ) has no impact on the probability of earning a reward once in the WZ, separated by offer (left) or collapsed across all offers (right, two-way ANOVA between group and OZ reaction time: $F_{2,9}$=1.657, $^{ns}p$=0.192). (c) The amount of time elapsed since the last pellet was earned has no impact on the probability of earning a reward once in the WZ, separated by offer presented upon arrival into the trial from which p(earn) is calculated (left) or collapsed across all offers (right, two-way ANOVA between group and time elapsed since last earn: $F_{2,9}$=1.601, $^{ns}p$=0.203). (d) The amount of time spent consuming and lingering at the reward site after earning a reward increases a function of time already waited during the countdown (i.e., offer length) split by ranked flavor preferences (left) or collapsed across restaurants (right). Significant main effect of time waited on post-consumption lingering time ($F_{29,39}$=17.704, *$p$<0.0001) but no significant interaction with group ($F_{2,29}$=0.549, $^{ns}p$=0.58). The data in (d) suggest mice are sensitive to time spent waiting in the wait zone, related to sensitivity to sunk costs during change-of-mind decisions. Error bars ± 1 SEM. Not significant (ns).

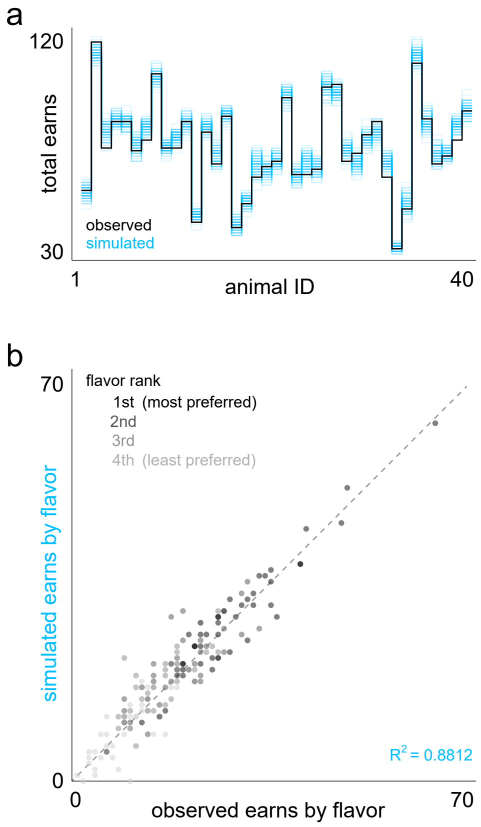


**Supplementary Figure 9. Economic model fit for the CREB cohort.** (a) Simulation of the Restaurant Row task quantifying total number of end-of-session earns, run 500 times (blue traces), in unique computer-generated sessions with randomly selected offers compared to observed earns (black trace) from an example session displayed across the 40 mice from the CREB cohort. (b) Scatter plot of observed and simulated earns split by restaurant ranked by subjective flavor preferences with a high degree of concordance.

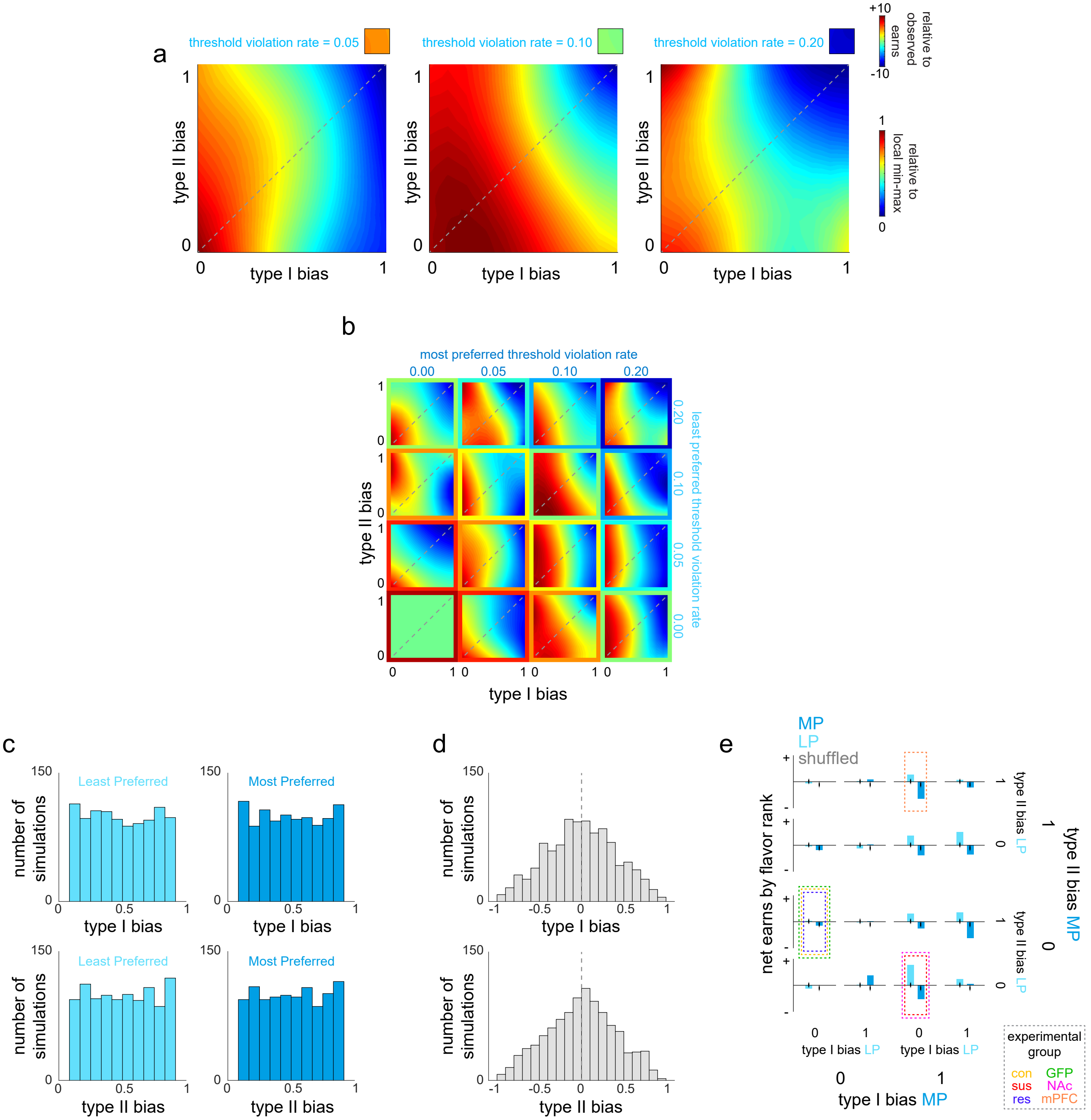

**Supplementary Figure 10. Economic model varying threshold violation rate and regret-related parameters.** (a) Simulations computing number of total pellets earned by varying the weight of type I violations on inflating the value of subsequent offers along the x-axis and the weight of type II violations on inflating the value of subsequent offers along the y-axis. Origin (0,0) indicates simulations ran where neither type of violation impacted subsequent decisions and individual trials were effectively treated as independent events. Coordinate (1,1) indicates simulations ran where both types of violations biased animals to always enter negatively valued offers on the subsequent trial. Violation rates set at 0.05 (left), 0.10 (middle), and 0.20 (right). Small inset colored-filled boxes to the right of each violation setting codes the difference in simulated number of total pellets earned relative to observed pellets earned. Low violation rate (0.05, left) yields higher than observed earns (orange-filled colored box) while high violation rate (0.20, right) yields lower than observed earns (blue-filled colored box). Medium 0.10 threshold violation rate (middle) yields a minimal difference relative to observed earns (green-filled colored box) and approximates observed violation rates. Color map of larger plots in panel (a) reflect relative number of total pellets earned normalized within each simulation, locally. (b) Violation rates were also systematically varied within each restaurant in these simulations, within the most preferred restaurant (across the top axis) and the least preferred restaurant (across the right axis) with interpolated violation rates for low and high preferred restaurants. The color of the border of each subplot indicates the relative difference in total pellets earned between simulated and observed data while the colormap of each subplot indicates the relative difference in total pellets earned normalized within each simulation locally. Note: the violation rate of 0.1 in both most and least preferred restaurants yield a subplot border color (green) that is closest to observed earns. Note: across all subplots, general trend of higher relative yield with lower type I bias regardless of level of type II bias persists. (c) Uniform distribution of bias settings used in 1,000 shuffled simulations with weights randomized independently in each restaurant. (d) Normal distribution of difference between bias weights between least and most preferred restaurant pairs in each shuffled simulation. (e) Simulations computing relative number of pellets earned split by the ranking of restaurants, displaying least preferred (LP, light blue) and most preferred (MP, dark blue) rewards earned. Simulations independently varied type I bias (bottom axis) and type II bias (right axis) each in LP and MP restaurants, binarized to 0 and 1 weights. The resulting matrix produced 16 economic phenotypes, only three of which were reminiscent of the social defeat and viral manipulation cohorts. Dashed boxes emphasized economic fingerprints characteristic of groups of mice from both the social defeat and CREB cohorts (color key defined in the legend inset). Horizontal line represents zero net change in relative rewards between simulated earns and average observed earns. Simulation categories were compared against 1,000 simulations randomly shuffling bias settings independently in each restaurant. Scale bar: ± 10 relative rewards. Note: the greatest change in net rewards earned by flavor stems from the combination of bias settings reminiscent of SUS and mCREB-NAc groups (type I bias: LP=0, MP=1; type II bias: LP=0, MP=0).
